# *Kita* crispants for systematic image-based genetic screens of complex traits in zebrafish larvae

**DOI:** 10.1101/2023.07.22.550161

**Authors:** Eugenia Mazzaferro, Christoph Metzendorf, Hanqing Zhang, Endrina Mujica, Ida Höijer, Ghazal Alavioon, Joao Campos Costa, Naomi L. Cook, Sara Gry Vienberg, Djordje Djordjevic, Anders Larsson, Adam Ameur, Anastasia Emmanouilidou, Amin Allalou, Marcel den Hoed

## Abstract

With thousands of loci identified by genome-wide association studies for complex traits, there is a need for *in vivo* model systems that can reliably and quickly infer the role of large numbers of candidate genes. CRISPR/Cas9-based functional screens in F_0_ zebrafish represent such a system. However, negative controls used so far – including scrambled guide RNAs (gRNAs), inactivated Cas9, and sham injections – do not elicit the same cellular and organismal responses as mutagenesis by CRISPR/Cas9, and may fuel biased conclusions. Here, we show that targeting *kita* facilitates efficient optical pre-screening for successful mutagenesis, higher quality imaging data, and efficient classification of cases and controls. We identified and tested two gRNAs that target *kita* with similarly high mutagenic efficiency and effects on pigmentation, and are free from off-target effects or major effects on cardiometabolic traits. We propose several approaches that will result in valid, unbiased conclusions.

## Introduction

In recent decades, genome-wide association studies (GWAS) have identified hundreds of loci that are robustly associated with complex diseases like type-2 diabetes (T2D) ^1^ and coronary artery disease ^2^. However, the clinical utility has so far been limited, because for the vast majority of loci, causal genes have not yet been functionally characterized. The biological relevance of genetic associations for human disease is traditionally investigated by perturbing candidate genes in cultured cells or murine models. Both model systems can provide valuable insights. However, while 2D and 3D cell cultures facilitate the required throughput, they lack the complexity of a live, intact vertebrate organism. Murine models on the other hand cannot be used to quickly and cost-efficiently examine the role in relevant traits for hundreds of candidate genes. Therefore, there is a need for *in vivo* model systems that can systematically characterize the role of a large number of genes in human diseases.

The zebrafish (*Danio rerio*) has numerous characteristics that make it a suitable model organism to functionally characterize the role of human genes. Zebrafish are vertebrate animals with a fully annotated genome with orthologues of ∼71% of human genes; or 82% of OMIM disease-associated genes ^3^. They have high fecundity, with female zebrafish laying >100 eggs per week. These eggs are fertilized externally, which makes them accessible for genetic perturbation using, e.g., CRISPR/Cas9. Early development is external and quick, with zebrafish embryos swimming with functional major organs by 48 hours post-fertilization. By 5 days post fertilization (dpf), all major organs are fully formed and functional. At the embryonic and early larval stages, zebrafish are optically transparent, allowing *in vivo* imaging of anatomical regions and biological processes using automated microscopy ^4^. Studies have shown that zebrafish larvae provide an attractive opportunity for *in vivo* genetic screens of cardiometabolic diseases, including T2D ^5^, fatty liver disease ^6^, and atherosclerosis ^7^.

CRISPR/Cas9 has dramatically simplified the targeted introduction of mutations in the zebrafish genome ^8^. Coupling Cas9 protein and a guide RNA (gRNA) into a ribonucleoprotein (RNP) complex can help introduce targeted double-stranded breaks virtually anywhere in the genome. The DNA damage triggers repair mechanisms that typically introduce indel mutations, which in turn disrupt protein function by introducing frameshifts and/or premature stop codons ^9^. Recent advances have made it possible to reliably generate biallelic knockouts directly in the microinjected embryos (i.e., in the F_0_ generation) ^10–12^. Previously reported genetic screens have demonstrated that F_0_ larvae can be used to examine effects on easily detectable monogenic morphological and behavioral traits ^10,12^, as well as on complex traits ^12–14^. Screening in these so-called crispants, without the need to generate and screen for homozygous or compound heterozygous mutants in the next generation, shortens the time frame of screens, and increases the chance to also identify relevant genes that reduce post-larval viability, fertility, and/or fecundity.

To reliably infer the role of a gene in disease etiology, it is desirable that crispants for the candidate gene only differ from sibling controls by a single factor. However, consensus has not been reached on the preferred controls. Some studies base their conclusions on a comparison of crispants with un-injected control larvae ^11,15^. However, a plethora of evidence from screens using morpholino oligonucleotides has firmly established that microinjections at the single-cell stage per se affect hatching, early development, and embryonic survival ^16–19^. Moreover, injections with Cas9 protein alone result in overexpression of genes involved in wound healing, already by 5 days post-fertilization (dpf) ^16^. Other studies generated controls by microinjecting Cas9 and (a) scrambled gRNA(s) ^10,12^, or by sham injections only ^20^. However, the absence of DNA editing in controls can be problematic, since DNA cleavage elicits an immune response ^19^. In yet other studies, controls were microinjected with a gRNA aimed at a control gene that was not targeted in cases for the candidate gene, e.g., the gene encoding tyrosinase (*tyr*) ^21^. In such cases, it is essential to ensure that on- and off-target effects of the control gRNA do not exert an effect on outcomes of interest. In the absence of epistasis, it is intuitive that cases and controls for the candidate gene should ideally both be targeted at the same control gene.

Control samples fundamentally influence the robustness of the results of any experimental pipeline. In this study, we aim to identify a gene that influences pigmentation, to facilitate pre-screening for successful mutagenesis and improved downstream image quality. Targeting the control gene should ideally not affect cardiometabolic traits, or induce CRISPR/Cas9 off-target effects elsewhere in the genome.

The primary role of *sparse*/Kit receptor tyrosine kinase-a (*kita*) in zebrafish is to regulate melanocyte specification, migration, and survival. *Kita* is expressed in melanoblasts and melanophores, and loss of its function prevents melanocyte migration, which ultimately results in apoptosis ^22,23^, rendering larvae free from pigment across the yolk and tail. Zebrafish also have a *kitb* gene, as well as genes encoding the *kitlga* and *kitlgb* ligands ^24^. Compromising *kita* and *kitlga* is sufficient to suppress melanophore migration and reduce pigmentation ^23^. In zebrafish, *kita* is expressed in the hematopoietic lineage, ovarian gametogenesis, and gastrointestinal tract, but *kita* null mutants have normal hematopoiesis ^25^, are fertile ^22^, and have no defects in digestion ^26^. This suggests its involvement in these pathways is not essential.

With the ever-increasing sample size of GWAS and improved ways to annotate associated regions, the list of candidate genes for cardiometabolic and other human diseases awaiting functional characterization is growing. Striving for an *in vivo* model system that can be used to functionally characterize genes in a systematic manner, we present a new, versatile control for CRISPR/Cas9- and image-based genetic screens in F_0_ zebrafish larvae. This approach will significantly reduce the screening time, improve image quality, and facilitate conclusions that are based on perturbation of the candidate gene, rather than on artifacts of the study design.

## Results

### Kita mutagenesis reduces pigmentation in 5-to-10-day-old zebrafish larvae

To identify a suitable control gene, we compiled a list of 13 zebrafish genes that affect pigmentation in larvae and for which human orthologues have not been associated with complex diseases. Based on a literature search, we excluded ten of these genes due to morphological, functional, or behavioral abnormalities in zebrafish (**Supp. Table 1**). To select the most appropriate of the three remaining genes (i.e., *kita*, *kitlga*, or *sox10*), we microinjected gRNA/Cas9 complexes with the highest predicted mutagenic efficiency into fertilized eggs, at the single-cell stage, and compared the effect on pigmentation (**Supp. Figures 1-2**, **Suppl. Table 2**).

All gRNAs successfully edit the DNA in the target site in >85% of microinjected larvae (**Suppl. Table 2**). At 10 dpf, *kita* mutagenized larvae display 3.5-fold less pigment (mean ± SD 4.9×10^4^ ± 2.9×10^4^ pixels) compared with un-injected siblings (1.7×10^5^ ± 4.2×10^4^ pixels, P_t-test_ = 3.3×10^-11^). This represents a larger effect than we observed for *sox10* (1.3- and 1.1-fold less pigment across the two gRNAs tested) and *kitlga* (1.9-fold less pigment). Moreover, all larvae that are mutagenized based on a PCR-based fragment length analysis (FLA) are free from pigment in the tail (<1.5×10^5^ pixels), compared with only 19% and 4% for the gRNAs targeting *sox10*, and 37% for *kitlga* (**Supp. Figures 3-4**). This indicates that the lack of pigment is most penetrant when targeting *kita*, and we therefore *pursued this gene as* the most promising control gene.

### Two kita gRNAs perform equally well and are free from off-target effects

To rule out the influence of environmental factors in a genetic screen, it is desirable that cases and controls for a candidate gene share environmental factors as much as possible. One way to accomplish this is by raising them in the same tank. In this approach, larvae eventually need to be classified as cases or controls. This can be done using a FLA for each targeted site in each candidate gene. However, optimizing primers and FLA conditions can become a rate limiting step. If *kita* is instead targeted at a different site in cases and controls, then classification may be possible by just genotyping larvae at the two *kita*-targeted sites, using an optimized FLA protocol. To examine if this is feasible, we next set out to identify a second gRNA, that targets *kita* with similar efficiency and effects.

The initially identified gRNA (*kita*-T1) targets the fourth exon of *kita*. We next identified a gRNA (*kita*-T2) that targets the second exon of *kita* (**Supp. Table 3**), with CRISPOR-predicted scores similar to *kita*-T1 (**Figure 1a**; **Supp. Table 3**). Both gRNAs target sites in the same immunoglobulin-like domain superfamily within *kita*. In embryos free from morphological defects at 1 dpf, 74.2% of larvae targeted with *kita*-T1 (95 of 128) and 89.4% of *kita*-T2-targeted larvae (59 of 66) survived until 5 dpf (**Supp. Table 3**). At 5 dpf, >95% of larvae targeted at *kita* exhibit less pigment in the tail compared with un-injected controls, irrespective of whether they were targeted at *kita*-T1 (96.8%) or *kita*-T2 (94.7%, **Figure 1c**). At 10 dpf, a FLA showed that 93% of larvae targeted using *kita*-T1 (40 of 43) have indel mutations in >80% of FLA peaks, compared with 98% of larvae targeted using *kita*-T2 (47 of 48, **Figure 1d**). Larvae with >80% of FLA peaks carrying indel mutations have ∼3-fold less pigment in the tail than un-injected sibling controls (P_t-test_=5.4×10^-11^), irrespective of the gRNA used to target *kita* (P_t-test_=0.7, **Figure 1b**). These results indicate that *kita*-T1 and *kita*-T2 perform similarly enough to allow differentiating between cases and controls for mutations in a candidate gene based on genotype at the *kita* targeted site.

**Figure 1.**
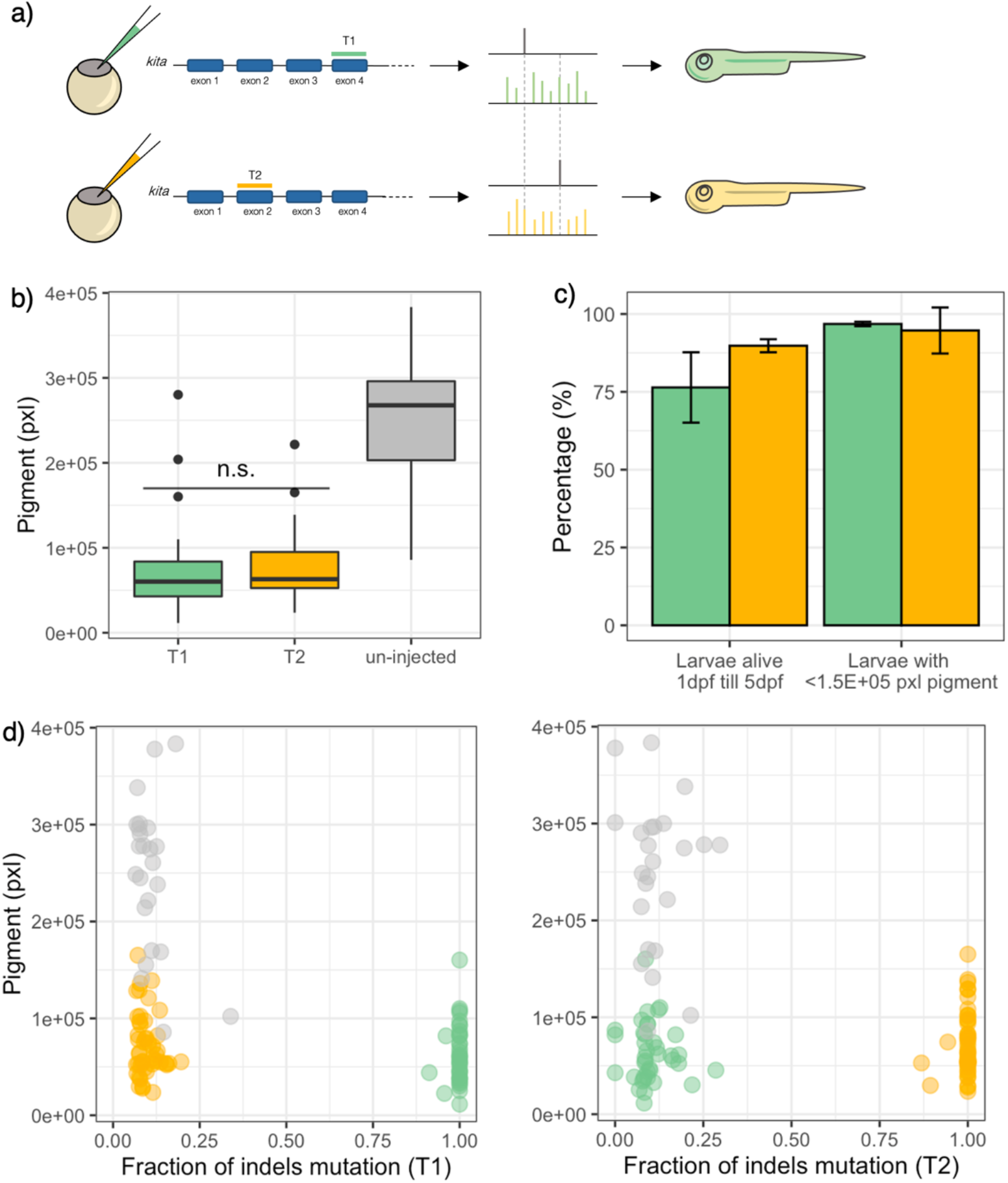
Targeting the *kita* gene in exon 2 (T2) and exon 4 (T1) with similar efficiency. **(a)** Microinjections at the single-cell stage to target *kita* in exon 4 (T1, green) or exon 2 (T2, yellow). **(b)** Boxplot to show the median level of pigment, i.e., the number of pigmented pixels in the tail, of larvae mutagenized with *kita*-T1 (n=44), *kita*-T2 (n=48), or un-injected sibling controls (n=28) at 10 dpf. One-way ANOVA test and Tukey’s Honest Significant Difference post-hoc test (P-value T1 vs T2 = 0.94). n.s., not significant**. (c)** Percentage of larvae targeted across four batches at T1 (n=128) or T2 (n=66) that were alive without a morphological defect at day 5; and the percentage of larvae with <1.5×10^5^ pixels of pigment in the tail. Bars represent percentages, error bars represent standard deviations. **(d)** Level of pigment in the tail (number of pixels) as a function of the fraction of total fragment length peaks that are indels, ranging from 0 (wildtype) to 1 (biallelic mutations in all cells).

### Both kita gRNAs are free from CRISPR/Cas9 off-target activity

A concern when using CRISPR/Cas9 is the potential to introduce mutations at unintended sites of the genome (i.e., off-target activity). We next examined if the two *kita* gRNAs have off-target activity that may result in undesirable and unanticipated mutations and phenotypic abnormalities. To account for the high genetic heterogeneity of zebrafish, and to accommodate for line-specific genetic differences, we evaluated off-target activity in DNA isolated from fin biopsies acquired from adult fish belonging to two different transgenic reporter backgrounds, one with fluorescently labelled beta cell nuclei and liver that we routinely use in our lab (i.e., Tg:-1.2*insH2B*:mCherry; 2.8*fabp10a*:GFP), and one with fluorescently labelled macrophages and neutrophils (Tg:*mpeg1*:mCherry; *mpo*:EGFP) ^27–31^ (**Supp. Figure 5**). In each transgenic reporter background – both are in the AB background – we extracted and pooled genomic DNA from biopsies of 10 adult fish. Using a genome-wide, amplification-free approach based on nanopore sequencing that we and others described earlier (Nano-OTS) ^32,33^, we only observed a clear double-stranded cleavage site at the on-target cut sites, and no other signals. This implies that both *kita*-T1 and *kita*-T2 are almost certainly free from off-target activity in the zebrafish we use in our experiments.

### Targeting kita exerts at most limited effects on cardiometabolic traits

Encouraged by the efficiency and specificity of *kita*-T1 and *kita*-T2, we next examined if targeting *kita* using either gRNA affects cardiometabolic traits (**Figure 2**, **Supp. Table 4**). Briefly, we used larvae from four different reporter backgrounds relevant for cardiometabolic studies. Siblings were either un-injected, microinjected with heat-inactivated Cas9, or microinjected with Cas9/*kita*-T1 or Cas9/*kita*-T2 (**Supp. Figure 6a**). Microinjection with either *kita* gRNA results in more deletion than insertion events (**Supp. Figure 6c**; **Supp. Figure 2**), and >95% of larvae targeted using *kita*-T1 and >97% of larvae targeted using *kita*-T2 have indel mutations in >80% of FLA peaks (**Supp. Figure 6d**; **Supp. Figure 2**). *Kita*-T1 may result in more unique mutations than *kita*-T2 (**Supp. Figure 6c**).

**Figure 2.**
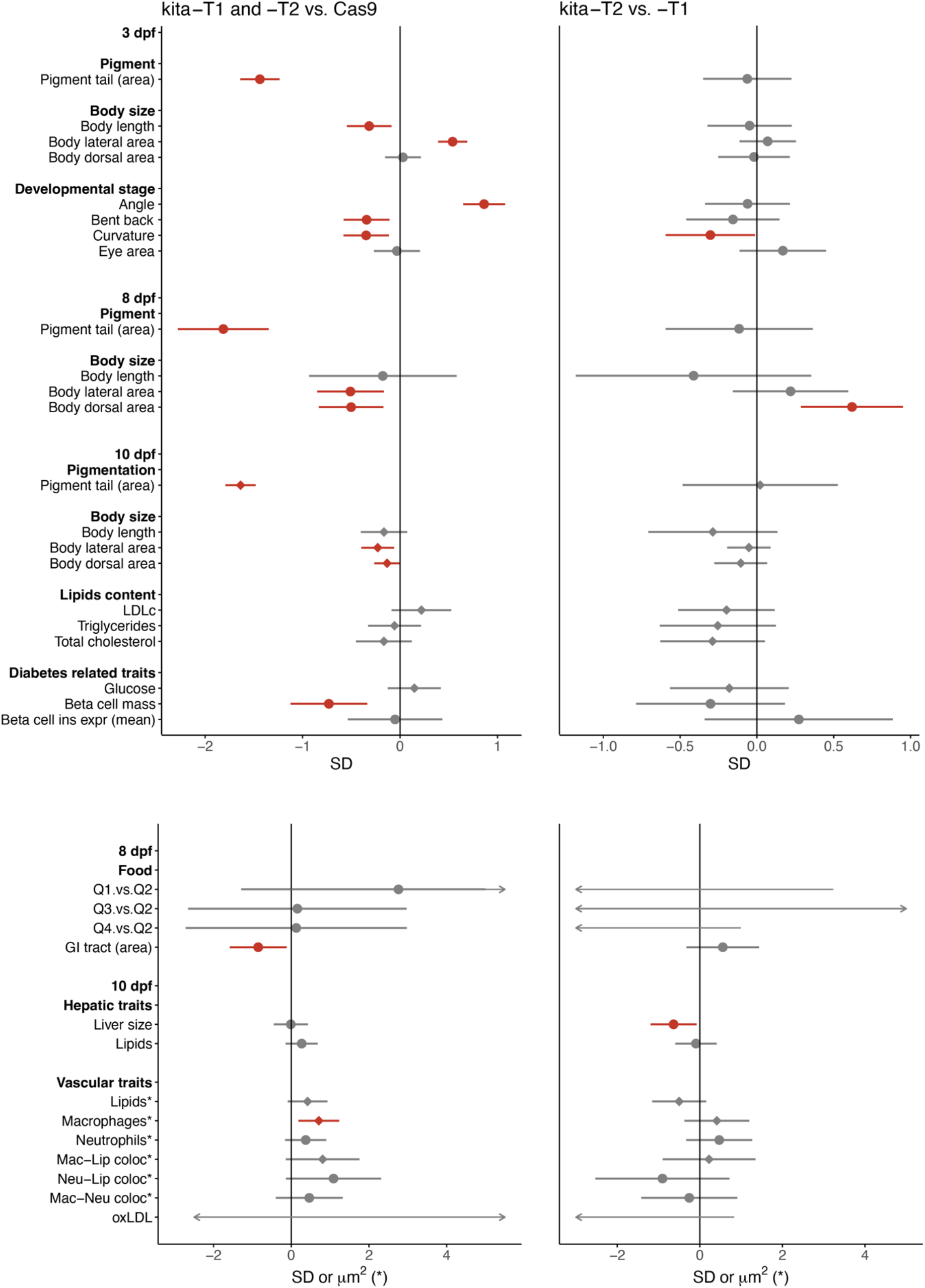
Effect of targeting *kita* using T1 or T2 compared with Cas9-only microinjected sibling controls on cardiometabolic traits in 3-, 8- and 10-day-old zebrafish larvae. Left: effect of targeting larvae at *kita* using T1 (n=94, 44, 108) or T2 (n=101, 46, 114) compared with microinjecting larvae with Cas9 protein only (n=111, 50, 173) from experiments in 3-, 8- and 10-day-old larvae, respectively. Right: effects of targeting using *kita*-T2 vs. *kita*-T1. Diamonds and dots show effect sizes; error bars show 95% confidence intervals of the effect size. Diamonds show the effect estimate for traits meta-analyzed across multiple transgenic reporter backgrounds (inverse variance weighted fixed effects meta-analysis); dots show the effect estimate based on the reporter background in which the trait was quantified. Effects shown in red have P < 0.05.

*Kita*-targeted larvae across all tested reporter lines (n up to 108 at *kita*-T1 and 114 at *kita*-T2) cluster together in terms of pigment and on average have 3.25-fold less pigment in the tail than the up to 173 sibling controls that were microinjected with Cas9 only (**Supp. Figure 6b**, **Figure 2**). Besides pigment in the tail, across the four reporter backgrounds, we examined seven traits reflecting early embryonic growth and development at 3 dpf; food intake at 8 dpf; and 17 traits reflecting body size; lipid and glucose content; pancreatic beta cell-related traits, hepatic traits; and vascular traits at 10 dpf. Compared with Cas9-only microinjected controls, *kita* mutagenized larvae have a delayed development at 3 dpf; and a smaller lateral and dorsal area normalized for body length at 10 dpf (beta ± SE -0.28 ± 0.09 and -0.21 ± 0.09 SD units, respectively, **Figure 2**, **Supp. Tables 5-6**). This may reflect a non-specific delay in development caused by DNA damage and repair per se. *Kita* mutagenized larvae also have a smaller pancreatic beta cell mass (-0.73 ± 0.20 SD units) – possibly reflecting a smaller body size – and more macrophages in the vessel wall (0.71 ± 0.27 SD units) than controls (**Supp. Table 5**). We cannot conclude if these effects are a direct result of mutations in *kita*, or if they reflect an immune response due to microinjection of gRNAs and Cas9 protein ^19^, DNA damage, and/or DNA repair.

We next examined if developmental and/or cardiometabolic traits are differentially affected by CRISPR/Cas9-induced mutations in *kita* at T2 vs. T1. Of all traits examined, mutations at *kita*-T2 result in shorter larvae at 10 dpf (-0.34 ± 0.16 SD units; n=176) with a smaller liver (-0.51 ± 0.21 SD units; n=115, **Figure 2**, **Supp. Tables 7-8**). Taken together, we do not observe strong evidence for effects of CRISPR/Cas9-induced mutations at either *kita* site (T1 or T2) on cardiometabolic traits at 3, 8 or 10 dpf.

### Targeting kita at two sites facilitates efficient genetic screens

To examine the suitability of *kita* as a control gene for efficient genetic screens in complex diseases, we examined if targeting cases and controls for candidates at *kita*-T1 and -T2 can be used to efficiently classify them for the presence or absence of mutations in candidate genes by means of genotyping at the *kita*-targeted sites. This method allows classification using a one-time optimized protocol for *kita*-genotyping, rather than having to optimize for each and every CRISPR/Cas9-targeted site instead. First, we examined if *kita* mutagenesis efficiency and reduction in pigment are affected by the co-injection of gRNAs against other sites. To this end, three gRNAs, targeting *kita*-T2 as well as exons 2 and 3 of a randomly selected candidate gene (*gpam*, a putative causal gene for triglyceride levels and liver fat ^30^ with a single orthologue in zebrafish), were microinjected together; while controls were microinjected with the gRNA targeting *kita*-T1 only (**Figure 3a**). Injecting one RNP (*kita*-T1) vs. three RNPs simultaneously (*kita*-T2, *gpam*-T1, and *gpam*-T2, **Supp. Table 9**) does not affect CRISPR efficiency (**Figure 3c**), the size of the indel mutations (**Supp. Figure 6c**) or the effect on pigment (**Figure 3d**).

**Figure 3.**
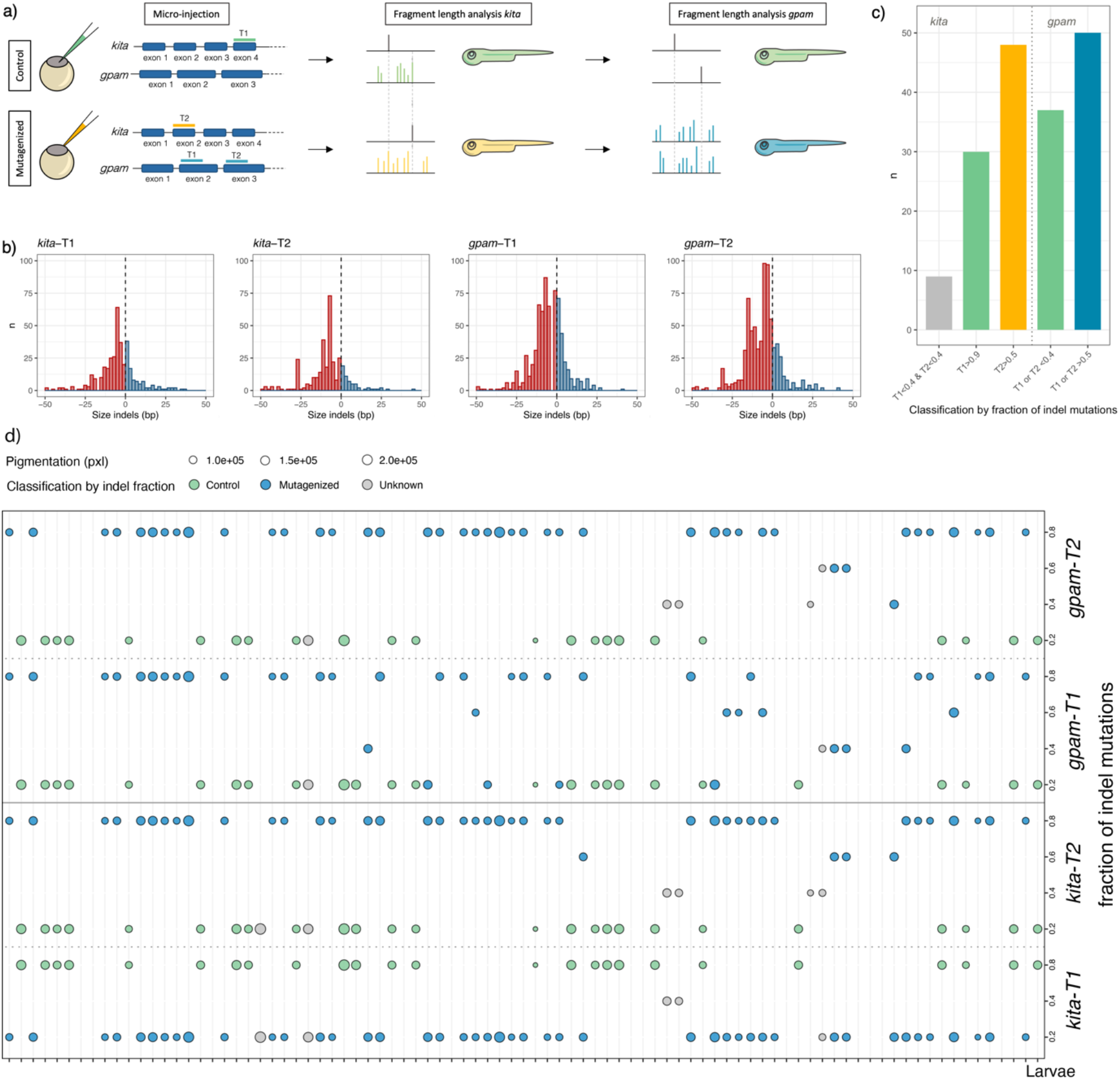
Targeting *kita* alongside a candidate gene. **(a)** Larvae were targeted at *kita*-T1 only, or at *kita*-T2 as well as at two sites in *gpam*. A PCR-based fragment length analysis was performed to assess the efficiency of the gRNAs targeting *kita* and *gpam*. **(b)** Frequency of each indel size (bp) for each of the gRNAs used: deletions are shown in red, insertions in blue. **(c)** Bar chart depicting the number of larvae assigned to be mutagenized (yellow or blue) or controls (green) based on a FLA of the regions flanking the *kita*-T1 (green), kita-T2 (yellow), or *gpam*-T1 and -T2 (blue) cut sites. **(d)** Dots showing – for each of the 87 larvae (x-axis) – the frequency of indels for each gRNA (y-axis). The color indicates if the larva was assigned to be mutagenized (blue), control (green), or undetermined (grey). The size of the dot represents the amount of pigment in the tail (smaller size, fewer pigmented pixels in the tail).

Next, we analyzed the efficiency of mutagenizing *kita*-T1 and -T2. Based on FLA genotyping results at *kita*-T1 (controls) and -T2 (*gpam* crispants), 48 larvae were classified as mutagenized at *gpam* (i.e., indels in >50% of FLA peaks at *kita*-T2 and <10% at *kita*-T1), and 30 larvae were classified as sibling controls (i.e., indels in >90% of FLA peaks at kita-T1 and <10% at kita-T2). Nine larvae (10%) could not be assigned to a group due to indel peaks in <40% of FLA peaks for both *kita*-T1 and *kita*-T2, indicating that mutagenesis in these larvae was inefficient, or the PCR quality was too low (**Figure 3b**). Genotyping based on amplification of the *gpam*-T1 and -T2 regions confirmed all calls made using FLA results for *kita*-T1 and -T2 for the 78 larvae. Of the nine larvae that could not be classified by genotyping *kita*, four were unmodified at both *gpam-*T1 and T2, while five were mutated at one of the two *gpam-*targeted sites.

Finally, we tested whether the mutagenesis rate at *kita*-T2 could be used as a proxy for the mutagenesis rate at *gpam.* Of the 43 larvae with indels in >80% of FLA peaks at *kita*- T2, 42 larvae were also successfully targeted (i.e., indels in >80% of FLA peaks) at *gpam*-T1 (n=1), -T2 (n=15), or both (n=26). The result shows that 97.7% of larvae that were highly affected at *kita*-T2 were also highly affected in the candidate gene (**Figure 3d**). This supports the hypothesis that genotyping by amplifying the regions flanking the cut-sites of *kita*-T1 and -T2 is sufficient to classify larvae as crispants or controls for the candidate gene. We next examined if the amount of pigment reflects the mutagenesis at *kita* as well as at *gpam*. Of the 75 larvae with a call at *kita* and data on pigment, 61 larvae have pigment-free tails. Four larvae (6%) had a pigment-free tail but at most moderate mutagenesis at *kita* (indels in <70% of FLA peaks at *kita*-T1 and -T2), of which only one was efficiently targeted at *gpam* (indels in >80% of FLA peaks at *gpam*-T1 or -T2, **Figure 3d**). This suggest that when cases and controls are raised in separate tanks, quantifying efficient CRISPR/Cas9-based mutagenesis at the candidate gene using pigment alone may lead to a 6% false positive rate. Moreover, only retaining less pigmented larvae on e.g., 5 dpf or the day of phenotyping can enrich the population for sufficiently mutagenized individuals, which can save time and reagents for genotyping.

### Cas9 and gRNA dosage effect

If controls are microinjected with a *kita* gRNA only, and cases are microinjected with a *kita* gRNA plus one or several gRNAs against a candidate gene, the effect that microinjection of different doses of RNP complexes (1 vs. 2 to 5) per se could have on the outcomes of the screen could be a concern. Others showed that the number of RNP complexes microinjected into eggs at the single-cell stage is associated with toxicity ^10^. To explore this further, we microinjected four sets of fertilized eggs with: i) RNP complexes of 1-5 gRNAs, each targeting a different site in *kita* (**Supp. Table 3**) at a 1:1 ratio of gRNA to Cas9 protein; ii) 5 gRNAs with 3 instead of 5 doses of Cas9 protein; iii) 1 gRNA with five doses of Cas9 protein; and iv) 1 dose of heat-inactivated Cas9 protein (**Figure 4a**). Survival to 5 dpf was low with microinjection of >3 doses of Cas9 protein. In larvae microinjected with 3 doses of Cas9 protein, survival was similar in those microinjected with 3 and 5 gRNAs (**Figure 4b, Supp. Table 10**), suggesting that high doses of Cas9 protein – and not of gRNAs – increase intracellular toxicity and reduce embryonic survival. This is in line with results reported by others ^10^.

**Figure 4.**
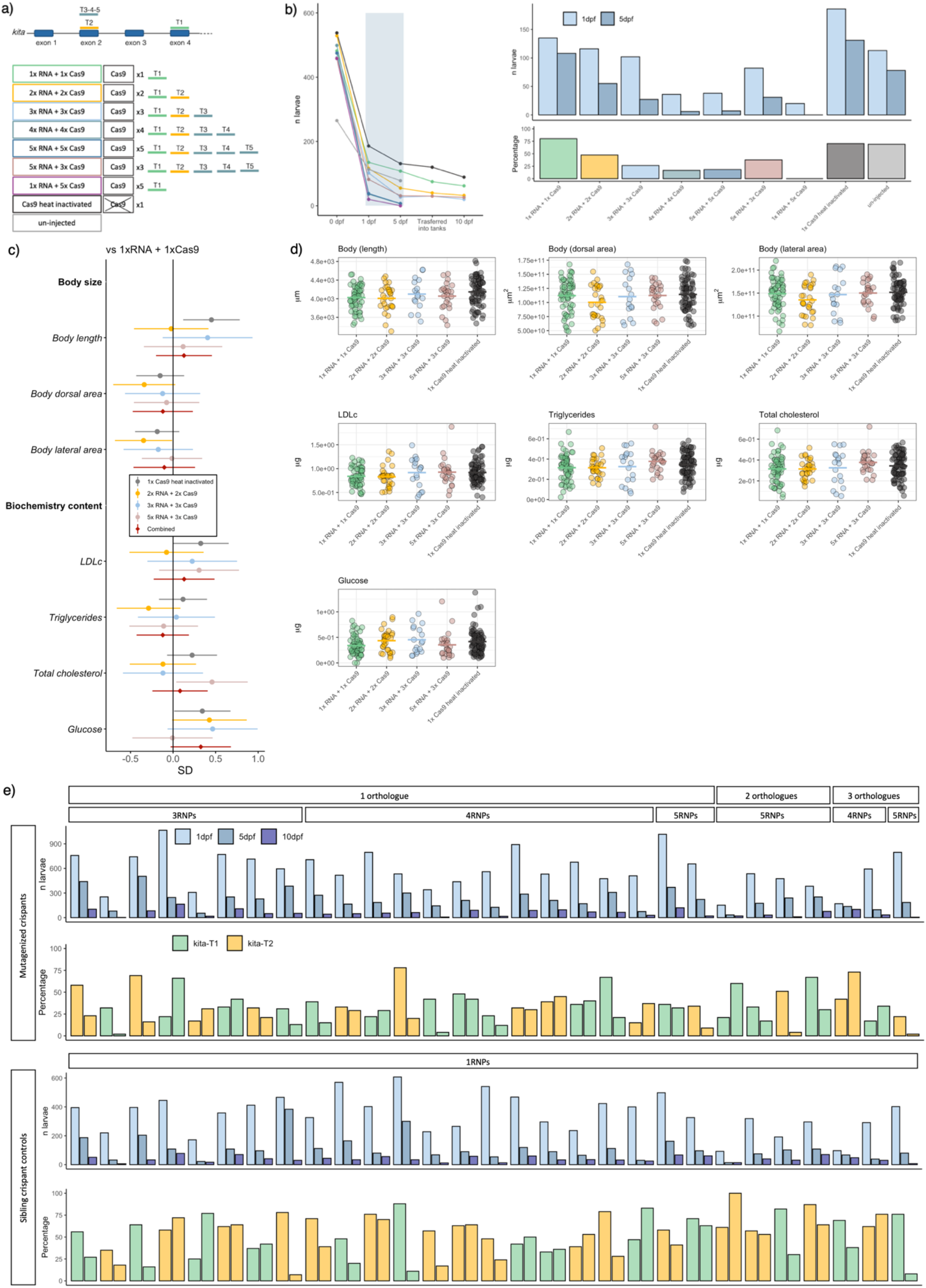
The effect of microinjecting multiple RNP complexes on viability and cardiometabolic traits. **(a)** Schematic overview of the *kita* exons targeted by gRNAs T1-T5. Larvae were microinjected with one of eight different combinations of numbers of gRNAs and accompanying doses of Cas9 protein. **(b)** The number of live larvae at 0, 1, 5, and 10 dpf (left) and the number (upper right) and percentage (lower right) of live larvae at 1 and 5 dpf for each of the eight conditions and for un-injected controls. **(c)** Diamonds and dots show effect sizes, error bars show 95% confidence intervals of the effect size. Dots show the effect of microinjecting 2, 3, or 5 vs. 1 gRNA complexes or of heat-inactivated Cas9 only (blue); diamonds show the joint effect of 2, 3 or 5 RNP complexes (red). **(d)** Distribution of raw values of the quantified traits for body size (upper) and whole-body cholesterol and glucose content (lower) in 10-day-old zebrafish larvae. **(e)** A total of 29 human genes with up to three zebrafish orthologues each were targeted using 3 to 5 RNPs per human candidate gene. Bar charts show – for the mutagenized crispants (top panel) and their *kita*-only targeted sibling controls (bottom panel) – the number of larvae that were alive at 1, 5, and 10 dpf (in blue), and the percentages of survival between 1 and 5 dpf, as well as between 5 and 10 dpf, with the colors indicating if the injection mix included *kita*-T1 (green) or *kita*-T2 (yellow). RNP: ribonucleoprotein.

To investigate effects on cardiometabolic traits, we raised larvae from conditions with at least 30 larvae surviving to 10 dpf and imaged them. Survival, pigment, body size, and whole-body LDLc, triglyceride, total cholesterol, and glucose content were quantified and larvae targeted at *kita*-T1 served as the control group (**Figure 4a**). Compared with 62 controls, the 26 larvae targeted using five gRNAs and three doses of Cas9 protein have 0.46 ± 0.21 SD units higher total cholesterol content, but are otherwise similar (**Supp. Table 11**, **Figure 4c-d**). The absence of a dose-response effect or of effects on other traits upon targeting *kita* at 2, 3 or 5 vs. 1 site suggests that within the studied range, these parameters are unlikely to be affected by the dose of microinjected gRNA and Cas9 protein.

Finally, we tested whether mutagenizing *kita* together with another gene using multiple RNPs increases mortality. We selected 29 human genes with 1-3 orthologues in zebrafish and simultaneously microinjected 3-5 RNPs. Here too, a larger number of microinjected RNPs results in lower survival from day 1 to day 5. However, we only observed <30% viability at days 5 and 10 when ≥5 RNPs (e.g., one RNPs targeting *kita* and ≥4 gene-specific RNPs) were co-injected (**Figure 4e; Suppl. Table 12**).

Taken together, our results suggest that targeting *kita* at different sites in cases and controls can facilitate more straightforward screens for human disease genes with up to three zebrafish orthologues.

## Discussion

Building on previous studies ^10–12^, we have developed a flexible, simple and fast method to functionally characterize genes for a role in complex diseases using zebrafish larvae. The exact approach used can vary depending on the needs of the screen. We were able to generate biallelic mutations in >85% of microinjected larvae in *kita* when targeting the gene using a single gRNA complex. Using FLA at just the *kita*-targeted sites, we could correctly assign >90% of phenotypically characterized larvae as cases or controls, with 97% congruence when compared with site-specific genotyping. By targeting cases and sibling controls at the same control gene, we ensure that all larvae are microinjected at the single-cell stage and undergo DNA editing and DNA repair. The two groups then differ by: 1) the amount of gRNA and Cas9 protein that is injected; 2) the site at which *kita* is targeted (optional); and 3) the presence of indel mutations in the candidate gene(s). The results we present here suggest that the first two differences are unlikely to materially influence a range of developmental and cardiometabolic traits.

We selected *kita* as our control gene, because mutations in this gene result in substantially less pigment in larvae of up to at least 10 dpf. The effects on developmental and cardiometabolic traits we observed were minor and may represent general effects of DNA damage and repair, rather than effects of targeting *kita* per se. Moreover, the effect might be due to the interference of melanocytes in the quantification of hepatic and vascular traits. Importantly, effects of mutations in *kita* on traits of interest are not problematic per se, since they are induced in cases as well as in controls for the candidate gene. Mouse models for atherosclerosis have suffered from such effects for decades, i.e., in apolipoprotein E (ApoE^−/−^) and LDL receptor (LDLr^−/−^) deficient mice ^35^. We anticipate that phenotypic consequences of mutations in *kita* are only problematic in the presence of epistasis with mutations simultaneously introduced in a candidate gene of interest. While epistatis cannot be ruled out for all candidate genes, it seems unlikely to be a systematic problem when aiming to functionally characterize the role of hundreds of candidate genes.

Studies in CRISPR/Cas9 founders require phenotypic characterization of microinjected animals, and results are affected by the success rate of those microinjections. In our experimental design, only larvae free from pigment at 5 dpf underwent further analysis, thereby removing larvae in which the microinjection and/or mutagenesis was not successful. A second advantage of this approach is the improved transparency of larvae, which benefits image-based phenotyping. Chemicals like 1-phenyl 2-thiourea (PTU) are routinely used to accomplish this ^36^. Our method provides a solution that avoids the need for such hazardous chemicals, which are arguably more likely to interact with mutations in candidate genes than targeted mutations in *kita* are. Moreover, we show that by image-based, automated quantification of pigment, we can distinguish between larvae that have been strongly mutagenized at *kita* and those that have not, reducing the need for assays to assess gRNA efficiency.

Earlier studies in the F_0_ generation suggested that the mutagenic effect of gRNAs should be verified before performing the actual experiment ^10–12^. Using the approach we propose, this pre-screening step can be replaced by evaluating the efficiency of the microinjection, mutagenesis, and DNA repair through quantification of pigment in the larvae that are actually included in the experiment. We show that the amount of pigment correlates highly with the activity of the gRNAs targeting *kita* and the candidate gene. One limitation is that drawing conclusions about mutagenesis in the candidate gene based on mutagenesis of *kita* or on pigment assumes the gRNA mutagenesis rate is high in *kita* as well as in the candidate gene. Based on our results upon targeting genes using up to four RNPs simultaneously, this is generally a safe assumption. Furthermore, others have shown that targeting a gene at multiple sites simultaneously increases the proportion of crispants with biallelic mutations, as compared with targeting a gene at one site only ^10–12^.

A possible disadvantage of functionally characterizing genes in the F_0_ generation is that the injected individuals are mosaic, i.e., they do not have uniform genotypes across all cells, and FLA does not specify the exact mutations present in each cell and cell type. Having a defined genotype could be advantageous if the aim is to study the effect of specific (human) mutations. However, when aiming to functionally characterize hundreds of genes using non-homologous end-joining, mosaicism is not problematic, as long as the mutation rate and proportion of frameshift mutations are high. In this scenario, characterizing mosaic larvae with a range of unique mutations within and across individuals, in several regions per gene, likely reduces the number of individuals in which compensatory mechanisms may still result in a functional protein. Importantly, the gRNA complexes induce editing within early stages of cell division ^11^, thereby increasing the probability that the tissue(s) of interest are affected, and reducing mosaicism as compared with earlier CRISPR/Cas9 protocols ^37^.

By co-injecting multiple RNP complexes, we increase the probability of double-stranded breaks in off-target sites. Recent studies using *in vitro* and synthetic methods have shown that off-target activity occurs more sporadically than previously anticipated ^33^. Another possible downside of this approach is reduced viability. Based on our results, performing experiments using the approach proposed here could be challenging for the ∼30% of human genes with multiple orthologues in zebrafish, since we show that microinjecting more than 3x the amount of Cas9 protein normally used to target a single site reduces viability by 5 dpf. However, we have successfully characterized human genes with two orthologues – confirmed by site-specific FLA – by maximally injecting up to 4x the normal dose of Cas9 protein and up to four gRNAs per microinjection.

To limit the scope for environmental factors influencing the results, it is generally beneficial to raise cases and control together, in the same experimental environment. In such cases, genotyping by a FLA at the *kita*-targeted sites is a quick and more efficient way to distinguish between cases and controls than gene-specific FLA at each CRISPR/Cas9-targeted site, since FLA optimization is typically required. In cases where genotyping might not be possible at all sites, using the *kita*-T1 and -T2 approach may also prevent loss of information. Alternatively, raising cases and sibling controls in separate tanks, with multiple tanks per condition could prevent the need for genotyping altogether. Tank-specific effects would become less likely to drive false positive genetic effects with more tanks per condition. This approach would further increase the throughput and scalability of genetic screens.

## Conclusions

We have developed a framework that facilitates the systematic functional characterization of genes for a role in complex diseases in crispant zebrafish larvae, by targeting mutagenized and sibling controls at *kita*. We identified two unique gRNAs that both generate larvae free from pigment, with biallelic disruption of *kita* in >84% of microinjected individuals. This facilitates *in vivo* enrichment of experimental larvae for successful crispants and efficient classification, likely without affecting development or cardiometabolic traits. This scalable method reduces experimental timeline from months to weeks or even hours – depending on the readout – and improves image quality.

## Methods

### Animal care

#### Zebrafish lines

Adult zebrafish were maintained at a temperature of 28°C in recirculating filtered water (Aquaneering, San Diego, CA), on a 14L:10D photoperiod with gradual dusk and dawn. Fish were fed twice per day with dry food (Sparos, Olhao, Portugal) and rotifers. Three transgenic reporter lines were generated by crossing transgenic zebrafish expressing fluorescent reporters on: 1) beta cell nuclei (Tg:-1.2*insH2b*:mCherry) ^30^ and hepatocytes (Tg:2.8*fabp10a*:GFP) ^27,31^; 2) macrophages (Tg:*mpeg1*:mCherry) ^28^ and neutrophils (Tg:*mpo*:GFP) ^29^; 3) macrophages (Tg:*mpeg1*:mCherry) ^28^ and oxidized LDL (Tg:*hsp70*:IK17-EGFP) ^38^. All transgenic reporter lines are maintained in the AB background.

#### Zebrafish experimental setup

The evening before microinjections, adult zebrafish were put into breeding tanks, with males and females separated by an optically clear plastic separator. The following morning, crossings were initiated by removing the separators and tilting an insert to generate a depth gradient. Eggs from different pairs were collected, mixed, briefly rinsed with clean water, and arranged on agarose stages for microinjections at room temperature (20-25 °C). Zebrafish embryos were incubated from 0 to 5 dpf at 28°C. On the morning of day 5, larvae that optically appeared free from pigment were transferred into 1 L tanks at a density of 30 larvae per 300 ml of filtered water. In the afternoon of day 5, larvae were first fed with 16.3 mg of dry food per feeding per tank of 30 larvae (Zebrafeed < 100 μm, Sparos, Portugal). The water in each tank was replaced by fresh water at 7 and 9 dpf, i.e., between the two feedings.

#### Microinjections

Microinjections of gRNA:Cas9 RNP complexes in fertilized eggs were performed as described by Hoshijima et al ^11^. Approximately 1 nL of the solution was injected into the cell of fertilized eggs at the single-cell stage. On the morning of 1 dpf, embryos were visually inspected to assess survival and to remove embryos with anatomical abnormalities. Microinjected embryos were incubated at 28.5 °C, in petri dishes with up to 60 embryos per dish. A subset of un-injected embryos from each injection batch was preserved until 5 dpf and then sacrificed for FLA using capillary electrophoresis.

### Guide RNA/Cas9 duplex preparation

#### Guide RNA design

For each of the three genes involved in pigmentation (*e.g*., *kita*, *kitlga*, and *sox10*), the community resource Zebrafish Information Network (ZFIN) was examined for proposed sequences of gRNAs that efficiently target early exons. The predicted efficiency was cross-checked using the CRISPOR v4.99 design tool ^39^. Guide RNAs were prioritized if they: 1) target early exons that are shared across transcripts; 2) have a high MIT specificity score; 3) have a high Cutting Frequency Determination score; 3) have a high Azimuth *in vitro* score; 4) have a high out of frame value; 5) have a high score for indels; and 6) have a low number of predicted off-target sites. If no gRNA was suggested by ZFIN, then gRNAs were designed manually using CRISPOR v4.99 ^39^, following the same criteria. The same approach was applied to design gRNAs for candidate genes associated with diseases.

#### Guide RNA and microinjection complex preparation

The Alt-R® CRISPR/Cas9 system from Integrated DNA Technologies (IDT, Coralville, Iowa, United States) was used as described elsewhere ^11,12^. Briefly, gRNAs made of a duplex of crRNA and tracrRNA (50μM) were prepared by mixing equal volumes of 100 μM Alt-R® crRNA and 100 μM Alt-R® tracrRNA in Duplex Buffer (IDT, Coralville, Iowa, United States) and annealing (95 °C for 5 min, gradual cooling at -0.1 °C/sec to 25 °C, and 25 °C for 5 min). gRNA:Cas9 RNP complexes were made by gently mixing 0.8 μL of 62 μM Cas9 protein stock (Alt-R® S.p. Cas9 nuclease, v.3, IDT, Coralville, Iowa, United States) for each 1 μL of 50 μM gRNA in a total volume of 14 μL. All gRNAs targeting orthologues of the same human gene were combined in a single solution. RNP complexes were incubated at 37 °C for 5 min and then kept at room temperature. Before injection, 1 μL of 0.5% Phenol red (ThermoFisher, Waltham, USA) was added to the solution to visualize micro-injections.

### Fragment length analysis for mutagenesis detection

#### Single gene amplification setup

Each larva or larval pellet after homogenization (e.g., mutagenized crispants, crispant controls, and 5-day old un-injected larvae) was incubated for 2 h at 55 °C and for 10 min at 95 °C in 50 μL lysis solution (proteinase K -Sigma-Aldrich, Munich, Germany) freshly diluted 1:100 in lysis buffer (25 mM NaOH, 0.2 mM EDTA), before pelleting the remaining debris by centrifugation at 3500 rpm and 4 °C for 3 min. The genetic region flanking the CRISPR/Cas9 cut site of a gRNA was amplified by PCR using forward primers with M13 priming sites at their 5’ end and pigtailed ^40^ reverse primer pairs (**Supp. Tables 2-3, 9**, IDT, Coralville, Iowa, United States). The reaction mix was assembled using OneTaq® DNA Polymerase (M0480) (New England Biolabs, Ipswich, MA, USA) and following the manufacturer’s instructions, i.e., 2 μL of lysed Genomic DNA was combined with 2 μL of 5X OneTaq Standard Reaction Buffer, 0.2 μL of 10 mM dNTPs, 0.2 μL of 10 μM of M13, 0.2 μL of primer mix (5 μM of forward primer and 10 μM of reverse primer), 0.05 μL of OneTaq DNA Polymerase, and water to a total volume of 10 μL. The following thermocycling profile was used: 94 °C for 30 sec; 34 cycles: 94 °C for 30 sec, 53-63 °C for 45 sec depending on primer pair, 68 °C for 30 sec; final extension at 69 °C for 5 min. The amplified products were diluted 5-fold with water, and 1.5 μL of sample was added to each well of a 96-well plate, which were pre-loaded with 10 μL of Hi-Di™ formamide (ThermoFisher, Waltham, USA) and GeneScan™ -400HD ROX™ Size standard (ThermoFisher, Waltham, USA) (9.85 μL HiDi formamide and 0.15 μL Size standard per well). To quantify the background signal of the FLA components, 8 wells did not receive samples. A further 8-16 wells were used to experimentally determine the size distribution of wild-type fragment sizes, using samples from un-injected (non-crispant) sibling control larvae. DNA was denatured by heating the plate to 95 °C for 5 min and rapidly cooling on ice, before FLA using an Applied Biosystems® 3730 DNA analyzer (Applied Biosystems).

#### Combined kita T1 and T2 gene amplification setup

The amplification of the genomic regions surrounding the two CRISPR/Cas9 cut sites in *kita*, referred to as T1 and T2, was performed in the same reaction as described for single gene amplification with the following modifications: a primer mix was assembled for each primer pair listed in Supp. Table 3 (IDT, Coralville, Iowa, United States). *Kita*-T1 primer mix: 18 μL non-fluorescent forward primer, 2 μL 5ATTO550N-fluorescent forward, 20 μL reverse primer, and water up to 200 μL. *kita*-T2 primer mix: 15 μL non-fluorescent forward primer, 5 μL 56-FAM-fluorescent forward primer, 20 μL reverse primer, and water up to 200 μL. Reactions were assembled as described above and the following changes were applied in the thermal profile: 30 cycles: 95 °C for 20 sec, 53 °C for 30 sec, and 68 °C for 30 sec. FLA was performed as described above.

#### Fragment length analysis: data analysis

Fragment sizes and peak heights were determined by analyzing the output data from the DNAnalyzer in Peak Scanner version 2.0 (ThermoFisher, Waltham, USA). Wild-type peak identification and relative quantification of mutant peak sizes per sample were determined using an in-house R v.4.1.0. script and Rstudio. Briefly, peaks of the forward primer’s fluorophore channel were selected, and background noise was determined from the peak heights in the 8 buffer-only wells. All peaks were discarded that were ≤95^th^ quantile of background peak height, as were all peaks with a fragment size outside the theoretical wild-type fragment size ±50 bp window, i.e., the size of most common indels caused by CRISPR/Cas9 mutagenesis ^41^. Peak sizes of all samples were rounded to full base pairs. The highest peak of 16 samples of un-injected sibling controls with a fragment size of theoretical wild-type fragment size ±1 base pair was selected as the wild-type peak. The median fragment size of the 16 samples of un-injected sibling controls was assigned as the wild-type fragment size.

Systematic differences between theoretical and experimental wild-type fragment sizes were corrected by adjusting for the difference between the theoretical size of wild-type fragments and the experimentally determined median fragment size of un-injected, wild-type control samples. Wild-type fragments in unknown samples were identified as corrected sample sizes of the theoretical wild-type size ±0.5 or a user-definable margin. If two peaks were within this size range, the peak closest to the theoretical wild-type fragment size was selected. Finally, all non-wild-type peaks were assigned to be in-frame or frameshift, depending on whether the difference in bp between the peak and the theoretical wild-type peak was a multiple of 3 or not. For each sample, the relative peak areas for the wild-type peak, frame-shift peaks, and non-frameshift peaks were subsequently calculated. Larvae were considered mutagenized if the relative wild-type peak area was <0.50, while larvae were considered sibling controls if they had a relative wild-type peak area >0.90. Larvae not fulfilling either criterion were excluded from the statistical analyses.

### Image acquisition

Larvae were cross sectionally phenotypically characterized for effects on early development at 3 dpf, food intake at 8 dpf, or for cardiometabolic outcomes at 10 dpf. For positioning and imaging whole-body morphology, we used the Vertebrate Automated Screening Technology (VAST) BioImager (Union Biometrica Inc., Geel, Belgium), which was mounted on the stage of a Leica DM6B automated fluorescence microscope with a Leica DFC9000 GT camera (MicroMedic A/B, Stockholm, Sweden). The fluorescence microscope was used to visualize transgenically labeled structures and organs.

#### Early embryonic development

In the morning of 3 dpf, larvae were transferred to clean water and anesthetized using 0.08 g/L tricaine solution (MS-222, Sigma, Sweden). They were then distributed into a 96-well plate at one larva per well, with 200 μL of tricaine solution per well. Each larva was automatically aspirated into a borosilicate glass capillary. Twelve tomographic images were acquired for each larva – one every 30° of rotation – using the brightfield camera of the VAST BioImager.

#### Food intake

Food intake was visualized and quantified in CRISPR/Cas9 founders from the AB background. Between 48 h to 24 h prior to imaging on 8 dpf, fluorescently labeled food was prepared by mixing 75 μL of yellow-green 2.0 μm polystyrene microspheres (FluoSpheres carboxylate-modified microspheres, Invitrogen, Carlsbad, CA, USA) with 54 mg of Zebrafeed, and 25 μL of deionized water. The mixture was dried at room temperature in the dark, and crushed into a fine powder on the morning of imaging.

In the morning of 8 dpf, batches of four larvae from each of eight tanks were transferred into 40 ml of clean water and fasted for 1 h, before being provided access to 2 μg of fluorescently labeled food per larva for 1 h. Next, larvae were transferred to clean water and anesthetized using 0.08 g/L tricaine solution (MS-222, Sigma, Sweden). Thirty-two larvae (4 larvae x 8 tanks) were then distributed into a 96-well plate at one larva per well, with 200 μL of tricaine solution per well. Each larva was then automatically aspirated into a borosilicate glass capillary. Twelve tomographic images were first acquired for each larva – one every 30° of rotation – using the brightfield camera of the VAST BioImager. Larvae were then positioned and oriented laterally to acquire two z-stacks of 30 optical sections (ΔZi, Zi-1 = 683.46μm) of the gastrointestinal tract using a 4X objective (Leica HC PL FLUOTAR 4X/0.13 DRY) at 10 msec exposures. One z-stack was in bright field and the other was obtained using the microscope’s L5 filter, to detect the fluorescently labeled food. To avoid bias by batch and time of day at imaging, we imaged larvae from each tank alternatingly and recorded the time of day at imaging. Seven rounds of imaging were performed to reach a total sample size of 204 larvae.

#### Pancreatic islet, hepatic, and vascular traits

The offspring from an in-cross of fish with fluorescently labeled beta cell nuclei and liver (Tg:-1.2*insH2b*:mCherry ^30^; 2.8*fabp10a*:GFP ^27,31^) were raised to 9 dpf as described above. In the morning of 10 dpf, batches of 4 larvae per tank were first transferred from their 1 L tank to 8 ml of fresh water. Next, larvae were transferred to a solution of 25 µM monodansylpentane (MDH) ^42,43^ in 1X PBS to stain lipid droplets. After incubation in the dark for 30 min, larvae were anesthetized, aspirated and imaged as described above (see: *Food intake image acquisition*). Here, larvae were positioned and oriented laterally to acquire a z-stack of 102 optical sections each (ΔZi, Zi-1 = 75.61μm) of pancreatic beta cells using the microscope’s TXR filter and a 40X objective (Leica HCX APO L U-V-I 0.80 W), at 20 msec exposures. After repositioning, three z-stacks of 75 optical sections each (ΔZi, Zi-1 = 151.75μm) were acquired of the bigger liver lobe using a 20X objective (Leica HCX APO L U-V-I 0.80 W). One z-stack was in bright field and acquired at 20 msec exposures. The hepatocytes were detected using the microscope’s L5 filter and the z-stack was acquired at 20 msec exposures. The MDH-stained hepatic lipid droplets were detected by the 405 filter at a 10 msec exposure.

Finally, the larvae were repositioned to acquire two z-stacks of 50 optical sections each (ΔZi, Zi-1 = 100.36μm) of the dorsal aorta and caudal vein in the proximal region of the tail, using the 20X objective. A bright field z-stack was acquired at 40 msec exposures; and the 405 filter was used to detect lipid accumulation in the vessel wall at 40 msec exposures. A total of eight batches were imaged to acquire a total sample size of 185 larvae.

#### Hepatic and vascular traits

CRISPR/Cas9 founders were generated in the offspring of an in-cross of fish with fluorescently labeled macrophages (Tg:*mpeg1*:mCherry) ^28^ and neutrophils (Tg:*mpo*:GFP) ^29^ and raised to day 9 as described above. In the morning of 10 dpf, batches of 4 larvae per tank were stained and anesthetized, positioned and imaged as described above (see: *Food intake*). First, two z-stacks of 70 optical sections (ΔZi, Zi-1 = 141.33μm) each were acquired in the liver region using the 20X objective. After repositioning, four more z-stacks of 50 optical sections each (ΔZi, Zi-1 = 100.36μm) were acquired in the tail region using the same objective. For both anatomical regions, the first z-stack was acquired in bright field; the others in fluorescence. The microscope’s TRX filter was used in the vasculature to visualize macrophages at 40 msec exposures; the L5 filter allowed visualizing neutrophils in the vasculature at 40 msec exposures; and the 405 filter detected lipid deposits in the liver at 15 msec exposures and in the vasculature at 10 msec exposures. A total of eight batches were imaged to acquire a final sample size of 180 larvae.

To visualize vascular accumulation of oxLDL at 10 dpf, tanks with CRISPR/Cas9 founders from an in-cross of fish with fluorescently labeled macrophages (Tg:*mpeg1*:mCherry) ^28^ and oxLDL (Tg:*hsp70*:IK17-EGFP) ^38^ were heat-shocked at 37 °C for 2 h in the morning of day 8, 15 min after the morning feeding. This procedure activates a transgenically expressed antibody (IK17) that specifically binds oxLDL ^38^. At 10 dpf, batches of 4 larvae per tank were stained and anesthetized as described above. This time, the L5 filter was used to visualize oxLDL, and the z-stack was acquired at a 40 msec exposure. A total of nine batches were imaged.

### Image analysis

The images acquired using the VAST BioImager’s camera (bright field) and the Leica DFC9000 GT camera (fluorescence) were automatically analyzed using a pipeline generated in-house that is based on a deep learning approach with data trained on thousands of images and run in Python and MATLAB 2022a.

#### Body size and pigmentation

The 12 tomographic images for each larva that were acquired using the VAST BioImager’s camera were analyzed using algorithms generated using PyTorch. Training sets with hundreds of manually segmented images were fed into the pipeline to recognize background, larva, and capillary. The algorithm next estimated body size parameters - i.e., body length, dorsal area, and lateral area - in images acquired at 3, 8 and 10 dpf. At each of these developmental stages, whole-body images of larvae in the dorsal orientation were used to quantify the amount of pigment in the tail. First, the algorithm segmented the region between the distal part of the swim bladder and the tip of the tail. Then, by applying thresholding, the melanocytes within this region were segmented. Finally, the area of the segmented region was calculated, providing an estimation of the amount of pigment in the tail of each larva. In whole-body bright field images acquired in 3 dpf larvae, images in the lateral orientation were used to segment and quantify the area of the eye, as well as to quantify the angle of the head and trunk, using the intercept of a line through the center of the eye and ear and one that follows the back contour of the larva.

#### Food intake

The fluorescence images acquired were deconvoluted and deblurred using the Leica software Thunder (MicroMedic A/B, Stockholm, Sweden). The z-stacks of optical sections from multiple focal depths in the bright field and fluorescence channels were combined into a single maximum intensity projection image. The maximal projection of the bright field images was used to segment the gastrointestinal tract. The maximal projection of the detected EGFP signal was used to quantify the fluorescently labeled food. Overlaying the two segmentations allowed quantification of the ingested food, i.e., the surface area of the fluorescence signal within the gastrointestinal tract.

#### Pancreatic beta cell nuclei

The optical sections of the z-stacks with detected mCherry-labelled beta cell nuclei were merged into a three-dimensional reconstruction of the pancreatic islet. From the 3D model, several parameters were estimated, including the number of beta cells, the total and average nuclear volume, and the average and total nuclear fluorescence intensity within each 3D object, as a proxy for beta cell insulin expression.

#### Liver

The frames from three z-stacks acquired for the liver region were processed into single maximum intensity projections, one for each channel. The maximal projection of the GFP channel (Tg:2.8*fabp10a*:GFP transgene ^27,31^) was used to segment the liver and to estimate its area in pixels. Lipid droplets were segmented as bright points from the maximal projection of the MDH ^42,43^ signal. By superimposing the two segmentations, the algorithm estimated the total number and area of lipid droplets in the liver.

#### Vascular traits

The two z-stacks were combined into a single maximum intensity projection. An algorithm was trained to recognize the circulation based on circulating lipids, and thereby recognize the ventro-caudal region of the tail, ranging from the lumen of the dorsal aorta to the ventral side of the caudal vein. As described above, the maximal projection in the MDH channel ^42,43^ was processed to segment the brightest spots as vascular lipid deposits. The combination of the two segmented images was used to quantify the surface area of lipid deposits localized in the region included between the caudal vein and the dorsal aorta. Earlier studies showed that these reflect lipid deposits in the vessel wall ^44^. Next, all the z-stacks in mCherry and EGFP were reduced to maximal projections by channel, and used to segment the total number and area of macrophages, neutrophils, or oxLDL, as well as the two-way co-localization between these traits and lipids.

#### Whole-body LDLc, triglyceride, total cholesterol, glucose, and protein content

After imaging, larvae were dispensed into the wells of 96-well plates. The anesthetized larvae were euthanized by exposure to 0.2 g/L tricaine (MS-222, Sigma, Sweden) on ice, and all excess water was removed before storing the sample at -20 °C. Each sample was subsequently submerged in 80 μL of ice-cold 1X PBS and homogenized (1600 MiniG-Automated homogenizer, Gammadata Instruments, Uppsala, Sweden) using two 1.4 mm zirconium beads (OPS Diagnostics, NJ, USA) for 2 min at 1,500 rpm. The samples were centrifuged for 5 min at 13,000 rpm at 4 °C, before 12.5 μL of sample was transferred into a 96-well, flat-bottomed plate with 12.5 μL of ice-cold 1X PBS and stored at -80°C, awaiting protein quantification. From the remaining supernatant, 70 μL was transferred to a 0.5 mL Eppendorf tube and stored at -80 °C, before quantifying LDLc, triglyceride, total cholesterol and glucose concentrations.

Protein quantification was performed using the Pierce bicinchoninic acid (BCA) Protein assay kit (ThermoFisher, Waltham, MA), at 562 nm using a Varioskan LUX Microplate Reader (ThermoFisher, Waltham, MA, USA), following the manufacturer’s instructions. LDLc, triglyceride, total cholesterol and glucose concentrations were quantified using direct LDLc (1E31), triglyceride (7D74), cholesterol (7D62), and glucose (3L82) reagents from Abbott Laboratories (Abbott Park, IL, USA) on a BS-380 chemistry analyzer (Mindray, Shenzhen, China). Concentrations were subsequently extrapolated to whole-larva quantities.

### CRISPR/Cas9 off-target activity

#### Tail fin transection

In each of the four reporter lines, 10 adult fish were anesthetized by immersion in ∼0.16 g/L tricaine solution and a partial transection of the tail fin was performed with a sterile scalpel. The tissue was transferred into Eppendorf tubes and stored in liquid nitrogen. The fish were immediately transferred into freshwater before being moved back into their original tanks. The tissues extracted from fish with the same transgenic background were pooled into a single Eppendorf tube and stored at -80 °C.

#### DNA extraction and purification

Genomic DNA was extracted from each pooled sample and purified using the MagAttract HMW DNA protocol (Qiagen, Hilden, Germany), with minor modifications. Briefly, 220 μL of ATL buffer was added to each Eppendorf tube and the tissue was disrupted with pestles, before adding 20 μL of Proteinase K. The samples were incubated at 56 °C at 900 rpm for 15 h. The DNA yield was quantified using a QUBIT kit 1x dsDNA HS Assay kit (ThermoFisher, Waltham, MA) and the quality was assessed using Femto Pulse at 1 mg/mL (Agilent, Santa Clara, CA).

#### Whole-genome long-read sequencing for off-target activity

Purified genomic DNA was sheared to 20-kb fragments by gTUBE (Covaris, Woburn, MA) by spinning the samples twice at 5,100 rpm for 60 sec. Samples were purified using AMPure XP beads, using 100 μL of freshly prepared 70% ethanol, and by eluting the sample with 25 μL of nuclease-free water to a concentration of 3 μg. The concentration of the DNA was measured using a QUBIT kit 1x dsDNA HS Assay kit (ThermoFisher, Waltham, MA) and fragment size was assessed on a Femto Pulse (Agilent, Santa Clara, CA). The sheared DNA samples were processed following V.3 of the Nano-OTS protocol ^32^. The long fragments were sequenced on a MinIONS (Nanoporetech, Oxford, United Kingdom) and reads were aligned using minimap2 ^45^. The presence of off-target activity was assessed using the custom-designed tool Insider ^32^.

### Statistical analysis

#### Within background analysis

We examined the effect of mutations on phenotypic traits in 3-, 8- and 10-day-old larvae within each of the appropriate reporter backgrounds. We only considered larvae mutagenized if >50% of their peaks area was of non-wild-type length and the larva had <1.5×10^5^ pigmented pixels in the tail region. Residuals – after adjusting for batch, tank and time of day at imaging – were normally distributed for most traits (**Supp. Figure 8**). Hepatic and vascular lipid deposition, and their co-localization with macrophages and neutrophils (examined for vascular lipids) show a negative binomial distribution, while food intake was enriched for zeros.

For variables with normally distributed residuals, we set observations to missing that were outside the mean ± 5xSD range. We next inverse-normally transformed normally distributed outcomes to a mean of zero and a standard deviation of 1, before examining the effect of exposures on cardiometabolic outcomes using linear regression analysis (regress in Stata). For the traits showing a negative binomial distribution, we used negative binomial regression (nbreg in Stata). Finally, effects of exposures on food intake were examined using multinomial logistic regression, with food intake divided into four quartiles of which the second quartile served as the reference group (mlogit in Stata). All analyses were adjusted for batch, tank and time of day at which the image was acquired.

#### Meta-analysis

For traits acquired in multiple backgrounds, we combined summary statistics across backgrounds using an inverse variable weighted fixed effects meta-analysis.

### Ethics

All experiments have been approved by the Uppsala Board for Animal Research (Djurförsöksetiska nämnd) of the Swedish ministry of agriculture (Jurdbruksverket, permit number Dnr 5.8.18-13680/2020).

### Data and code availability

Only standard code was used in this project. All summary statistics are made available as supplementary tables.

## Acknowledgements

Computations were performed on resources provided by the Swedish National Infrastructure for Computing (SNIC) through the Uppsala Multidisciplinary Center for Advanced Computational Science (UPPMAX) under project SNIC 2022/22-235. Handling and storage of imaging data were enabled by resources provided by SNIC, partially funded by the Swedish Research Council through grant agreement no. 2018-05973. Nanopore sequencing was performed by the SciLifeLab National Genomics Infrastructure (NGI) in Uppsala, Sweden.

## Supplementary Figures Index

**Supplementary Figure 1.**
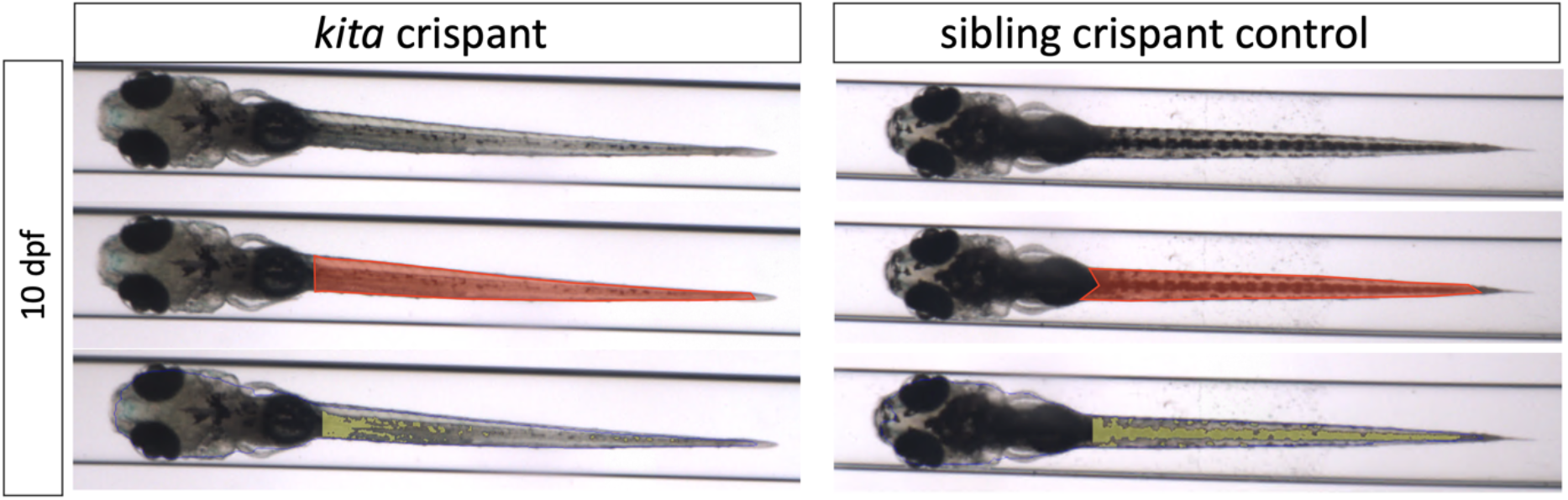
Effect of targeting *kita* at T1 or T2 on pigment in the tail. Dorsal view of a *kita*-T1 or -T2 targeted zebrafish larva with an average level of pigment (for the condition) in the tail (left) and an un-injected sibling control (right) at 10 dpf. Quantification of pigment in the tail was performed by segmenting the tail between the distal part of the swim bladder and the tip of the tail (red segmentation, middle panel) from the raw images (top panel). A threshold was applied to segment the pigmented area (yellow segmentation) in the segmented tail region (bottom panel).

**Supplementary Figure 2.**
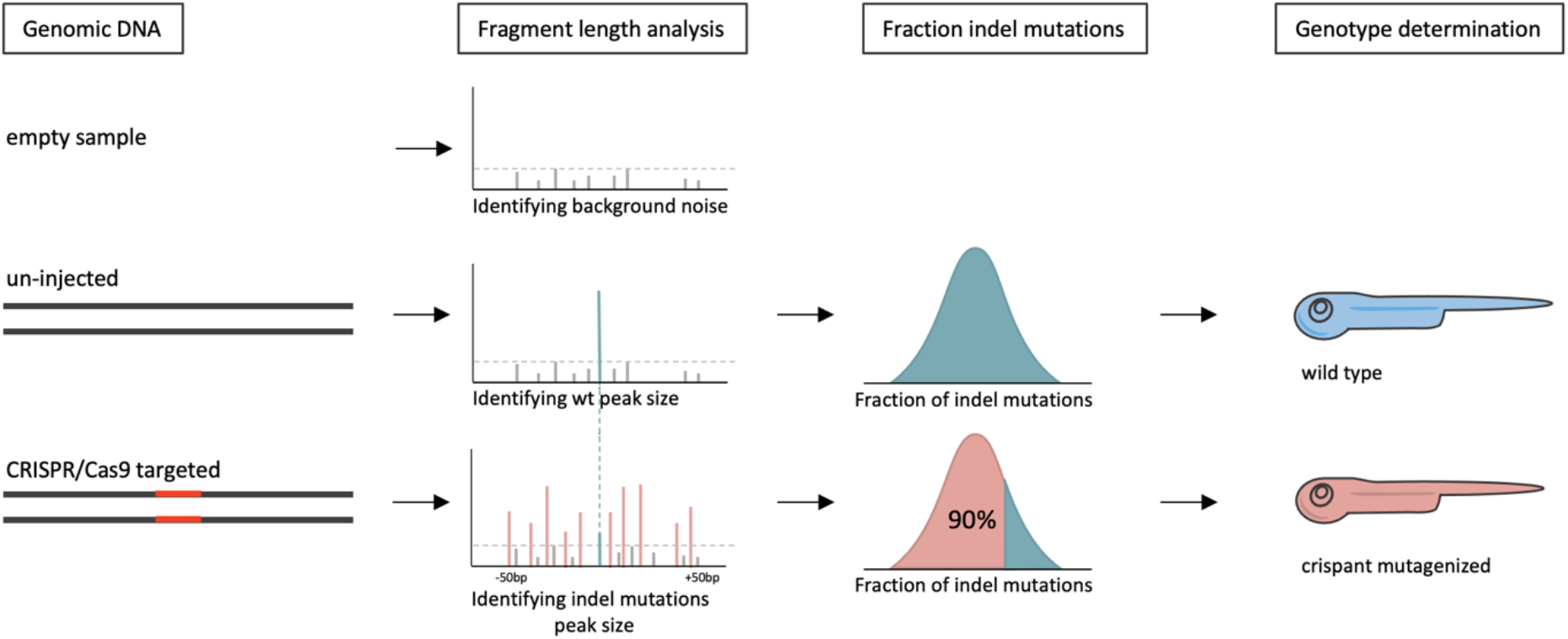
The main steps to identify mutagenized larvae directly using analysis of PCR product. The target region is amplified with fluorescence-labelled primers; capillary electrophoresis separates the fragments from background noise (i.e., from empty samples) and quantifies the height of the wild-type peak (from un-injected siblings) and indels within ±50 bp of the CRISPR/Cas9 cut site. For each larva, the fraction of total peak area made up of indel mutations is computed to distinguish mutagenized larvae (fraction >0.50) from sibling controls (fraction <0.10).

**Supplementary Figure 3.**
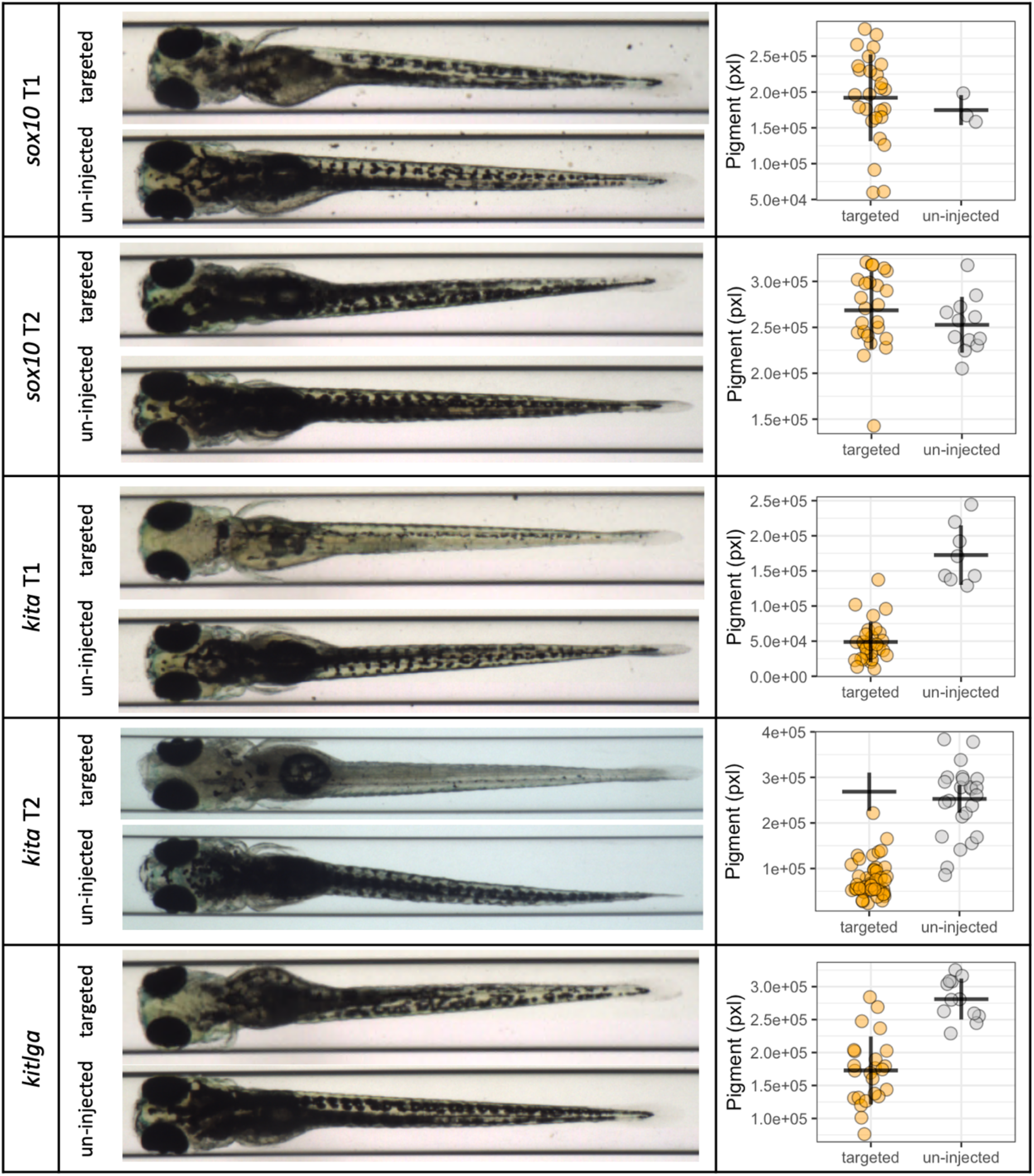
Pigment in the tail of 5-day-old zebrafish larvae targeted at three genes known to affect pigmentation. Dorsal view of average – for the condition – 5-day-old zebrafish larvae targeted using *sox10*-T1 or -T2, *kita*-T1 or -T2, or *kitlga*-T1. The amount of pigment was compared between CRISPR/Cas9-mutagenized individuals (orange) and un-injected sibling controls (grey). Plots show the amount of pigment in the tail in targeted larvae and in un-injected sibling controls. Lines show average values and SD.

**Supplementary Figure 4.**
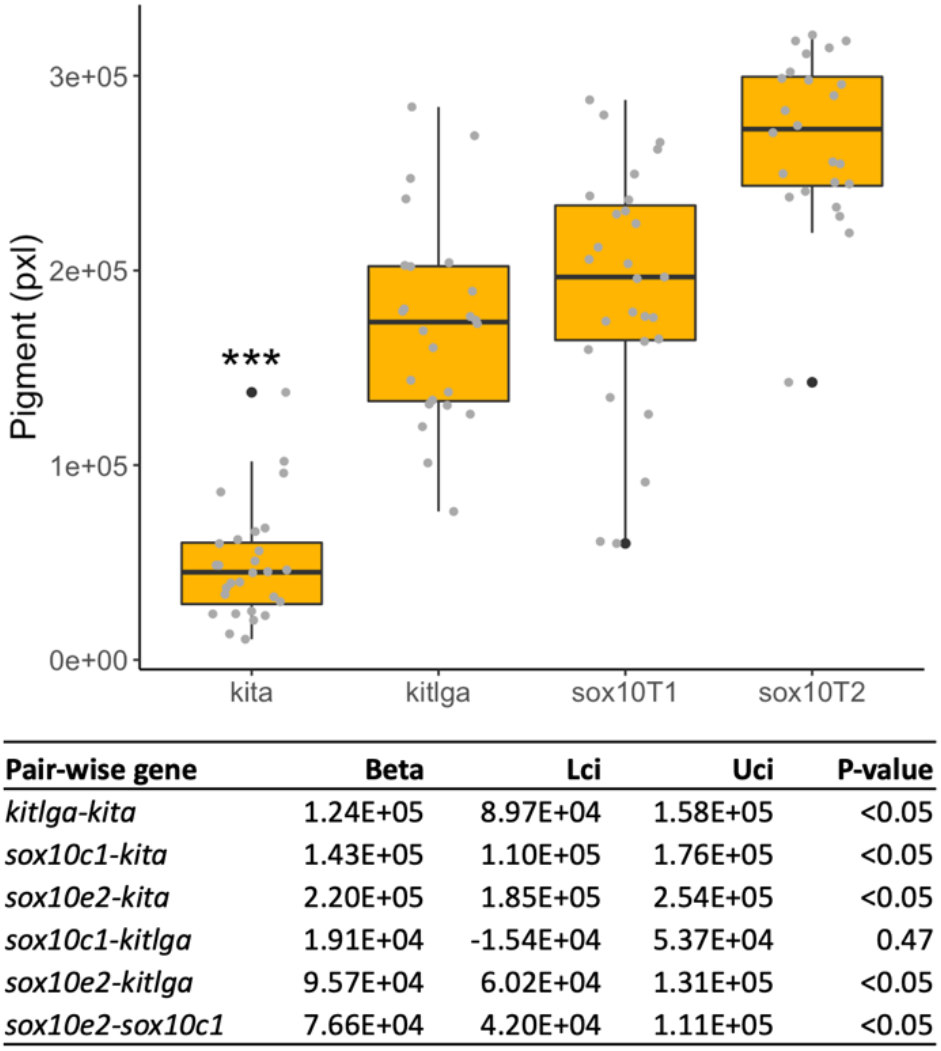
Pigment in the tail of 10-day-old zebrafish larvae targeted at one of three genes known to affect pigmentation. Box plot showing the amount of pigment in the tail (in pixels) for zebrafish larvae targeted at *kita*, *kitlga*, and *sox10*. The table shows one-way ANOVA results for pair-wise comparison of targeted larvae based on fragment length analysis and un-injected controls. Lci and Uci: lower and upper 95% confidence intervals.

**Supplementary Figure 5.**
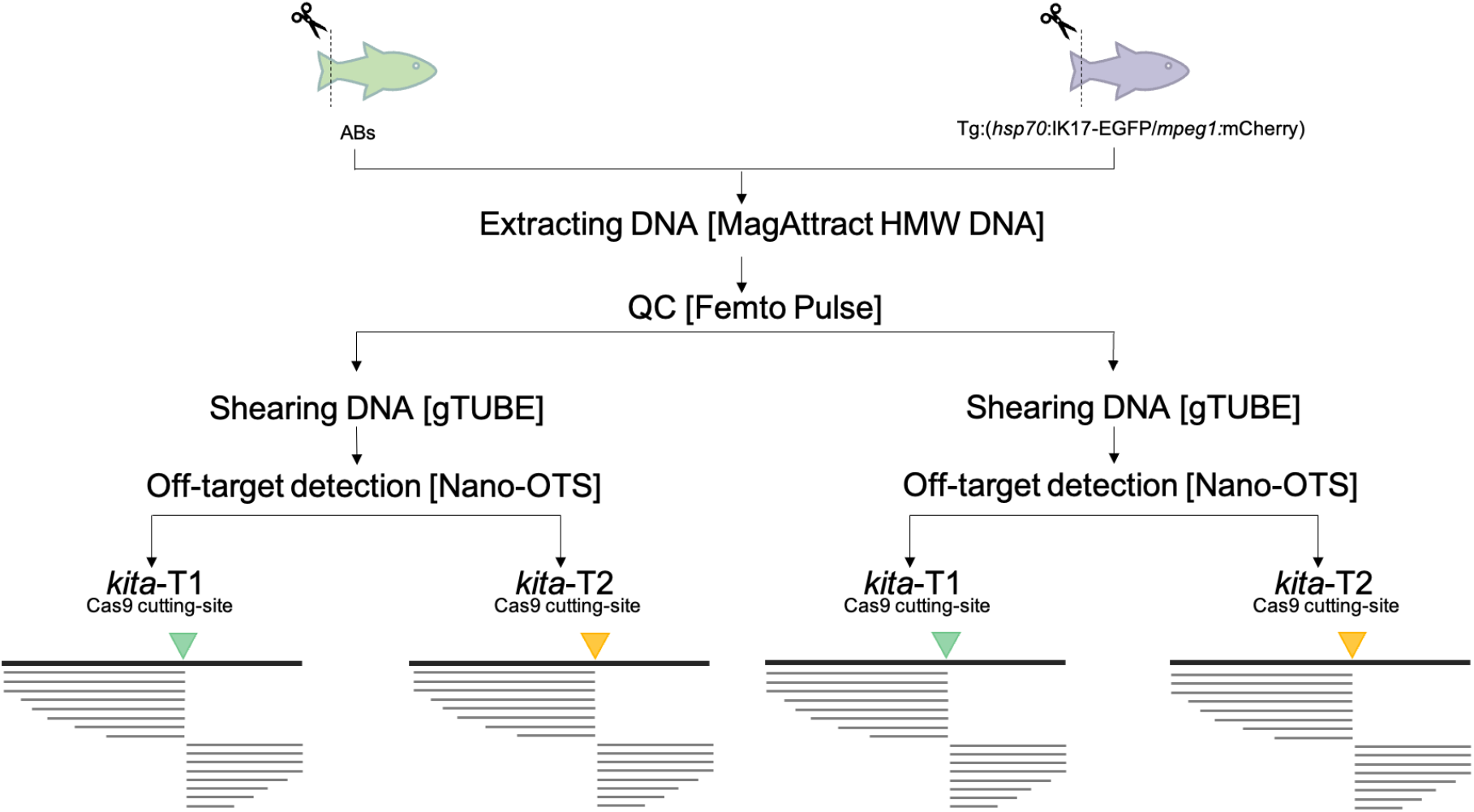
Detecting potential off-target activity for *kita*-T1 and *kita*-T2 duplex gRNAs. Tail fin transections were collected from two different reporter lines – both in the AB background – at 10 individual adult fish per reporter line. The genomic DNA was extracted and purified using the MagAttract High Weight Molecule DNA; the size fragments were detected by Femto Pulse. The samples were sheared to 20-kb fragments by gTUBE before being processed on MinIONS following the Nano-Off-Target Sequencing v.3 protocol. The identified Cas9 cut sites are the ones anticipated for T1 and T2.

**Supplementary Figure 6.**
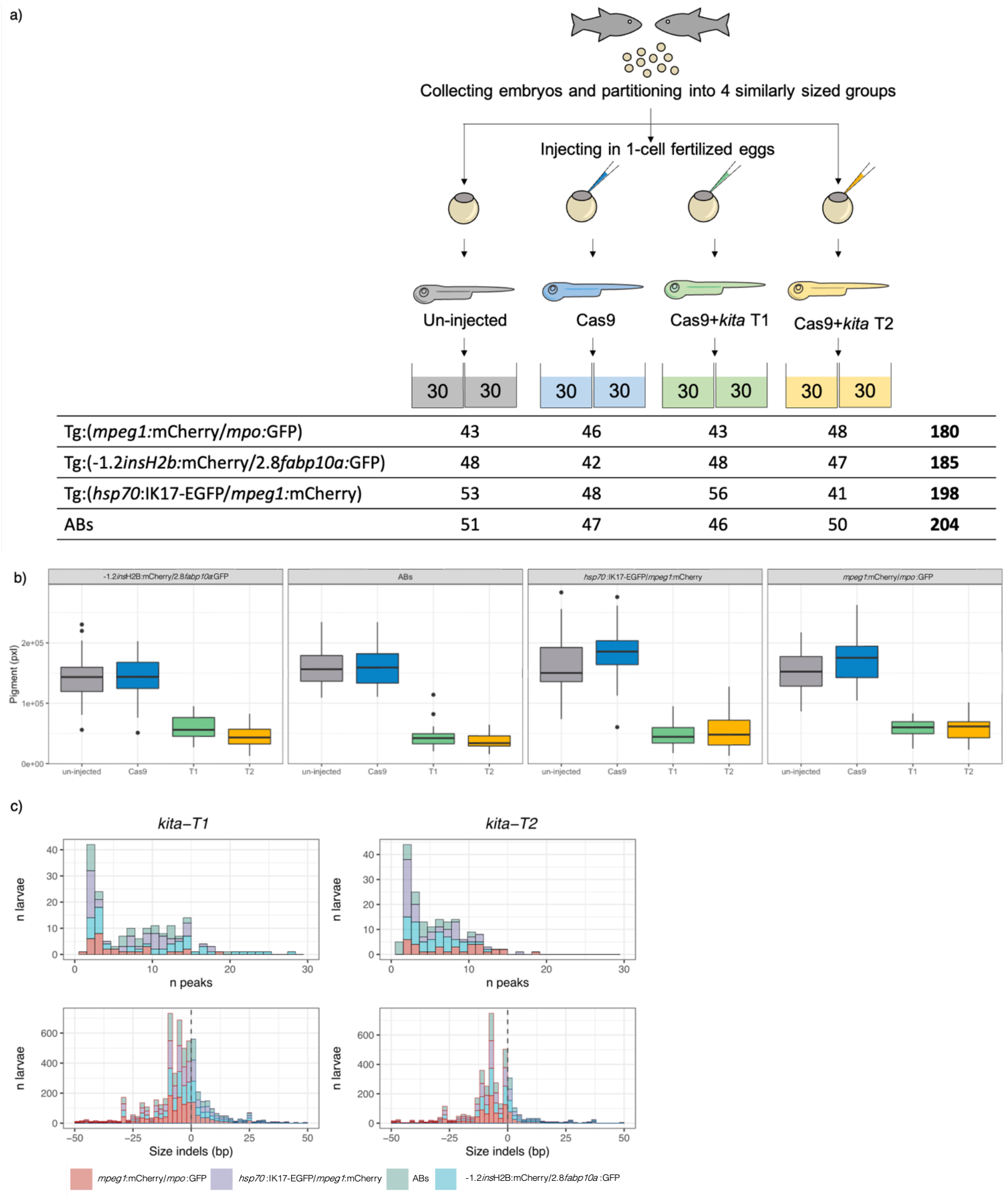
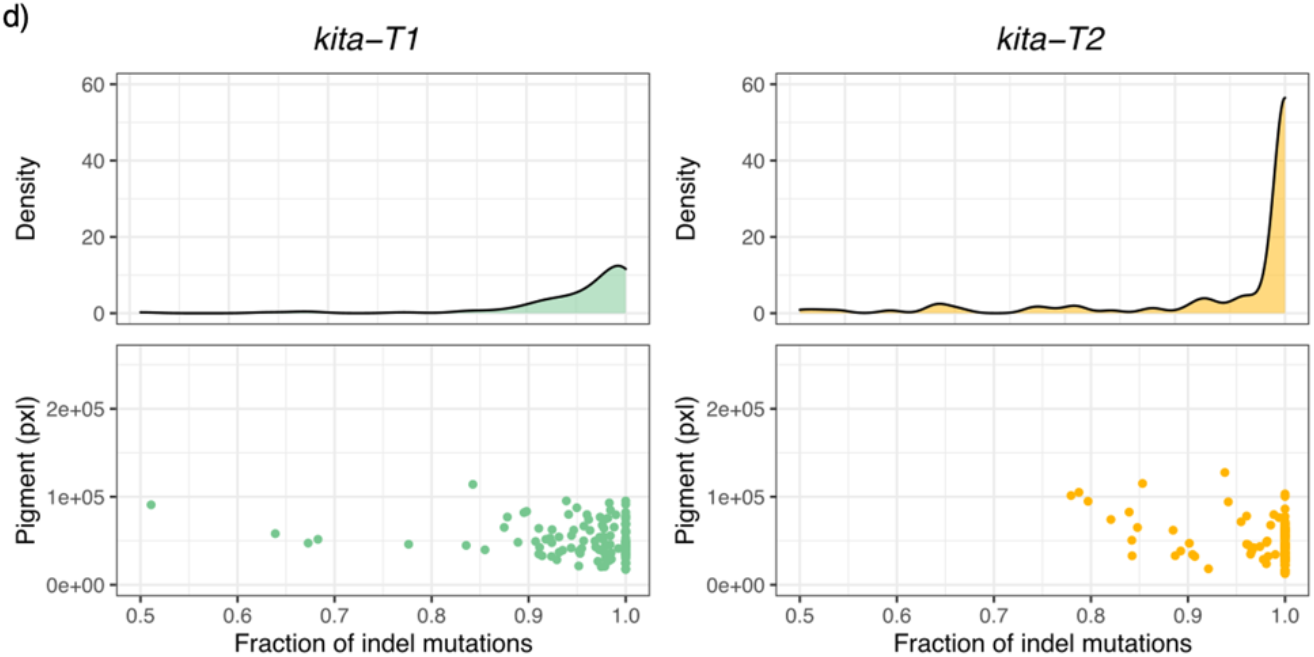
Effect of targeting *kita* using the T1 or T2 duplex gRNAs on pigment in 10-day-old zebrafish larvae. **(a)** After an in-cross of zebrafish that are homozygous for the reporter transgene(s), eggs were collected and randomly divided into four equally sized groups. Group 1 was left un-injected (grey); group 2 was microinjected at the single cell stage with only Cas9 protein (blue); the other two groups were targeted at the *kita* gene using duplexes with either the T1 (green) or the T2 gRNA (yellow). At day 5, 60 larvae from each group were transferred into two tanks (30 larvae/tank) and fed twice daily until they were characterized for traits facilitated by their fluorescently labelled transgenes at day 8 or 10. The table reports the number of imaged larvae in each of the four reporter lines. **(b)** Distribution of the amount of pigment in the tail, stratified by condition. **(c)** The number of indel mutation peaks per larva (upper) and the frequency of indel size (bp) within ±50 bp of the cut site (lower) of *kita*-T1 (left) and *kita*-T2 (right). Deletions are shown to the left of the dotted line; insertions to the right of the dotted line. **(d)** Density curve of number of larvae targeted in *kita* using T1 (left) and T2 (right) as a function of the fraction of total FLA peaks that are indel mutations (upper). Scatter plot of pigmented pixels in the tail as a function of the fraction of total FLA peaks that are indel mutations (lower).

**Supplementary Figure 7.**
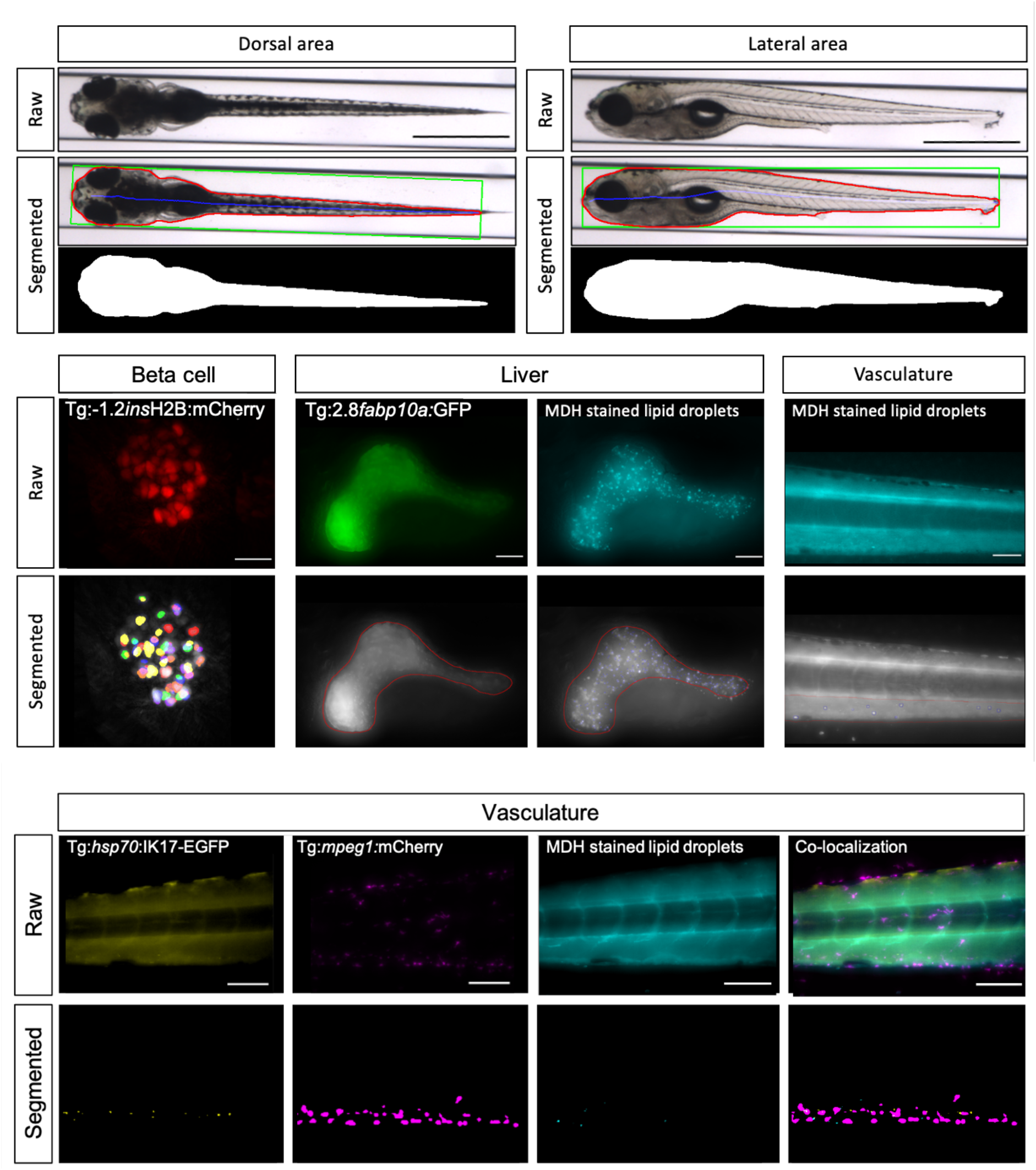
Visualizing and automatically segmenting whole-body and tissue-specific traits in 10-day-old zebrafish larvae. Top: dorsal and lateral view of 10-day old zebrafish larva and segmentation of length (length green line in dorsal view) and area. Middle: visualizing and segmenting of 1) beta cell nuclei (Tg:-1.2*ins*H2B:mCherry) to quantify beta cell mass (total nuclear volume); and average and total beta cell nuclear insulin expression (i.e., fluorescence intensity in segmented objects, scale bar=10μm); 2) lateral view of fluorescently labelled hepatocytes (Tg:2.8*fabp10a:*GFP) for liver size; 3) hepatic lipid deposition (monodansylpentane, MDH) (scale bar 100μm); and 4) vascular lipid accumulation (monodansylpentane, MDH, scale bar 100μm). Bottom: visualizing and segmenting vascular accumulation of a) oxidized LDL (Tg:*hsp70*:IK17-EGFP); b) macrophages (Tg:*mpeg1*:mCherry); c) lipids (monodansylpentane, MDH) and d) their co-localization (scale bar 100μm).

**Supplementary Figure 8.**
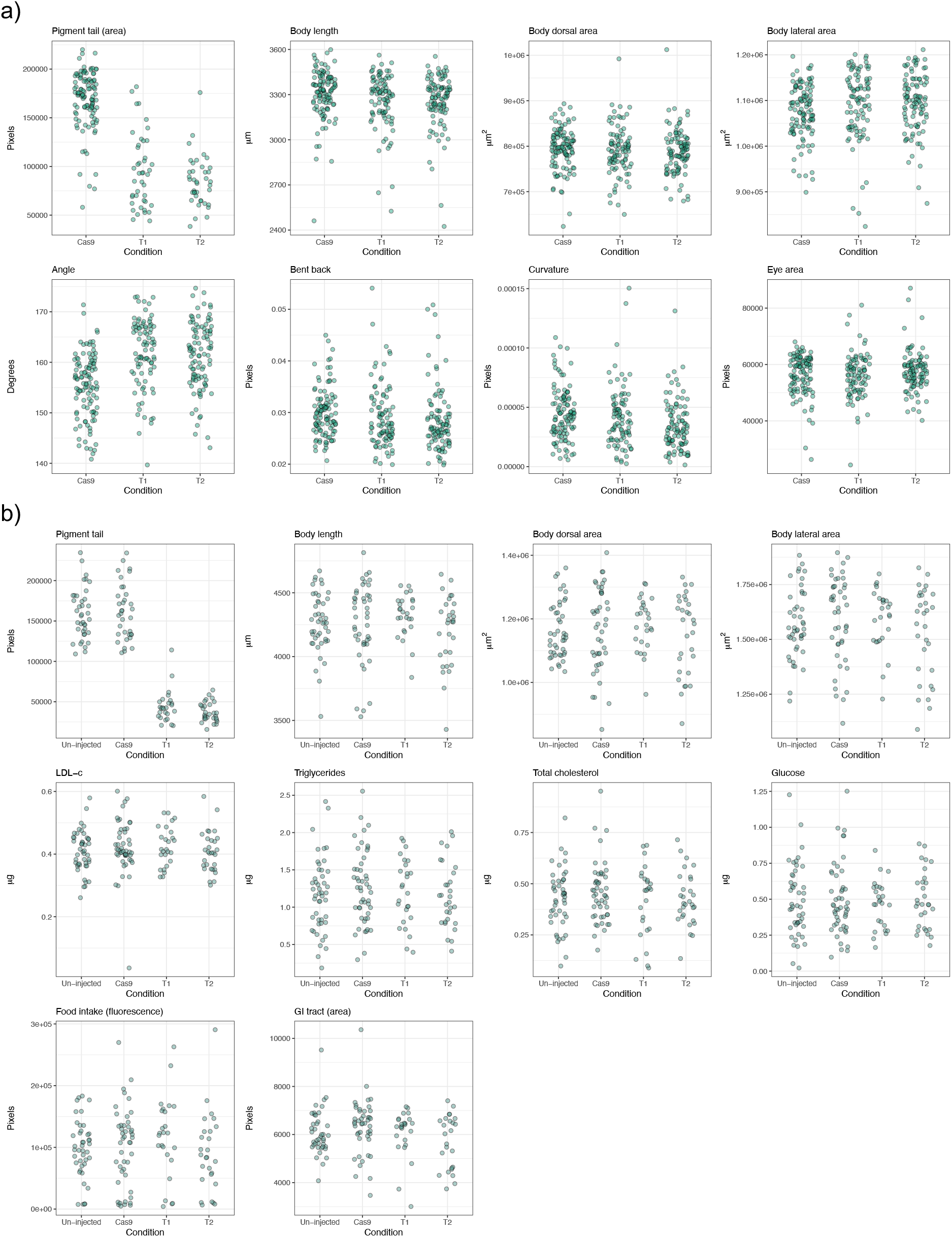
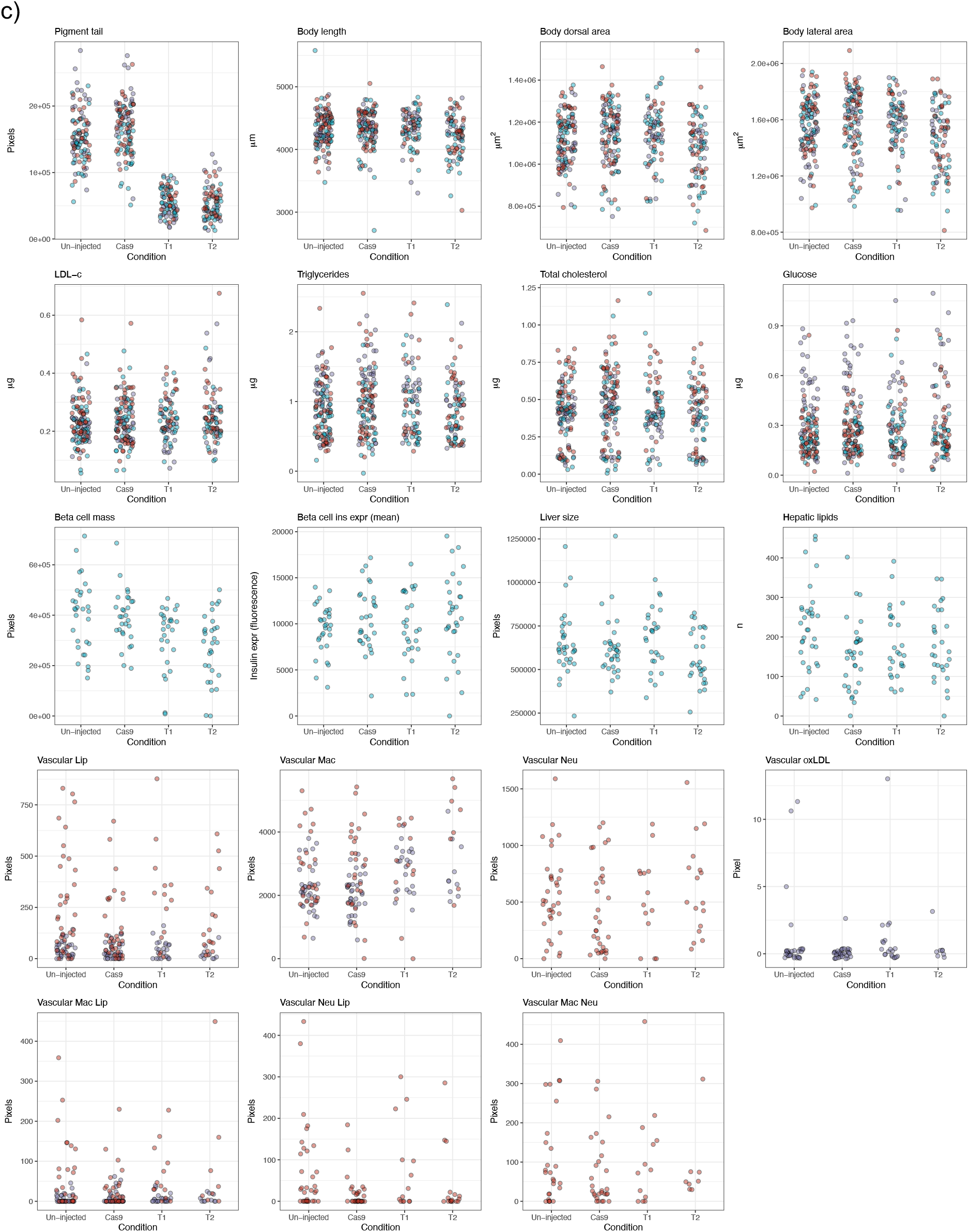
Raw traits quantified in 3-, 8-, and 10-day-old zebrafish larvae. Pigment in the tail and body size traits; food intake traits; whole-body cholesterol and glucose-related traits; pancreatic beta cell traits; liver traits; vascular traits. Colors depict the background: light blue: Tg:(-1.2*ins*H2b:mCherry / 2.8*fabp10a*:GFP); purple: Tg:(*hsp70*:IK17-EGFP / *mpeg1*:mCherry); red: Tg:(*mpeg1*:mCherry / *mpo*:GFP); green: ABs.

